# First Order Associations Between Banff Acute Lesions in Kidney Allograft Biopsies and a Urinary Cell Three-Gene Diagnostic Signature

**DOI:** 10.64898/2026.07.22.740171

**Authors:** Carol Li, Joseph E. Schwartz, Thalia Salinas, Darshana M. Dadhania, Alex DeVito, William Higgins, Steven Salvatore, Surya V. Seshan, Vijay K. Sharma, Thangamani Muthukumar, Manikkam Suthanthiran

## Abstract

Banff acute lesion scores underpin histologic classification of kidney allograft biopsies; however, biomarker studies rely on second-order associations with diagnostic categories that introduce confounding. We quantified the first-order relationships between Banff acute lesion scores and the validated urinary cell three-gene rejection signature. In 354 biopsy-urine pairs, three-gene signature scores computed using a locked regression equation incorporating absolute copy numbers of CD3E mRNA, CXCL10 mRNA, and 18S rRNA in urinary cell RNA-were related to glomerulitis (*g*), peritubular capillaritis (*ptc*), interstitial inflammation (*i*), and tubulitis (*t*). Signature scores rose monotonically with Banff acute lesion severity, with 1.5 to 1.8-fold higher odds of more severed *g, ptc, i,* and *t* (all P<0.0001), and showed good calibration. Associations remained robust for composite microvascular (*g+ptc*) and tubulointerstitial (*i+t*) indices and were strongest for severe *g* and *t*, supporting this signature as a noninvasive, quantitative readout of acute rejection pathology with immediate diagnostic applicability.

---

Kidney transplantation confers substantial gains in survival and quality of life compared with dialysis for patients with end-stage kidney disease, ^1–4^ but these advantages are undermined by acute rejection, a major driver of allograft loss. Thus, early, accurate, and mechanistically informative detection of acute rejection remains central to preventing irreversible damage.

The Banff classification is the global reference standard for diagnosing and categorizing kidney allograft rejection ^5–7^. Central to this framework are the Banff acute lesion scores, which semi-quantitatively grade histopathological changes in glomerulitis (*g*), peritubular capillaritis (*ptc*), interstitial inflammation (*i*), tubulitis (*t*), and intimal arteritis (v) on a scale from 0 to 3, providing a granular readout of rejection activity ^8,9^. Recent work has fundamentally reshaped the use of these scores: continuous or composite lesion-based indices consistently stratify allograft failure risk more effectively than traditional categorical Banff diagnoses, such as antibody mediated rejection (ABMR) ^10^. A Banff-based histologic chronicity index ^11^ and a related prognostic index ^12^ derived from chronic lesion scores in patients with ABMR showed a strong, independent association with graft loss and provided a simplified tool for clinical decision-making. Moreover, an automated Banff-driven histological decision-support system that relies on lesion scores reclassified more than 40% of biopsies relative to original pathologist-assigned diagnostic categories, and this reclassification significantly improved prediction of long-term allograft outcomes ^13^. Also, analyses of about 19,500 kidney transplant biopsies demonstrated that continuous indices constructed from Banff lesion scores capture the full spectrum of rejection activity and chronicity, outperform categorical Banff diagnoses in discriminating rejection phenotypes and independently predicting graft failure ^14^. Together, these advances support a paradigm shift: from exclusive reliance on Banff diagnostic categories toward systematic incorporation of lesion scores and lesion-based indices as primary descriptors of intragraft biology in both clinical decision-making and risk stratification ^15^. However, lesion scoring requires an invasive biopsy with attendant complications ^16–18^. The field therefore urgently needs noninvasive biomarkers that can read out the same granular rejection biology captured by Banff lesion scores without tissue sampling.

In the Clinical Trials in Organ Transplantation (CTOT)-04 study, we developed and validated a urinary cell three-gene signature comprising CD3E mRNA, CXCL10 (IP-10) mRNA and 18S rRNA levels (urinary cell three-gene signature) that was diagnostic and prognostic for acute cellular rejection ^19^. This signature achieved an area under the curve of 0.85 for distinguishing acute cellular rejection from non-rejection in a large, prospective cohort of 4,300 urine specimens from 485 kidney allograft recipients.

Banff diagnostic categories are defined by algorithmic combinations of multiple Banff lesion scores, and these composite rules have been repeatedly refined over successive Banff updates ^5–9,20–22^. Biomarker associations with these composite diagnostic categories (second order) may simply reflect the underlying individual lesion scores (first order) that define them. In this context, first-order associations—direct correlations between biomarkers and individual Banff lesion scores—offer a methodological advantage analogous to the endophenotype strategy in psychiatric genetics, where mechanistically narrower component traits yield cleaner biological associations than heterogeneous composite diagnoses ^23^. Contemporary biomarker evaluation frameworks emphasize analytical rigor, transparency of interpretation and minimization of confounding ^24,25^. Despite this conceptual appeal, first-order biomarker-lesion relationships in kidney transplantation have rarely been examined systematically. CTOT-04 and subsequent other biomarker studies have almost uniformly evaluated second-order associations—correlating biomarkers with composite biopsy diagnoses such as TCMR or ABMR—rather than directly with the lesion scores that drive those diagnoses ^19,26–30^.

Here, we address this gap by reframing the CTOT-04 urinary cell three-gene signature as a noninvasive, quantitative reporter of Banff lesion biology, rather than of categorical rejection diagnoses. We hypothesized that the three-gene signature would demonstrate robust associations with Banff acute lesion scores for *g*, *ptc*, *i*, and *t*, and that these relationships would clarify which components of intragraft inflammation are most strongly encoded in the urinary cell signature. We analyzed 354 biopsy-matched urine specimens from kidney allograft recipients, correlating the three-gene signature with individual Banff lesion scores and composite lesion combinations using contingency analyses, and show that the urinary cell three-gene signature is tightly coupled to the severity of acute Banff lesions, particularly tubulitis and glomerulitis, quantitatively tracks composite microvascular and tubulointerstitial inflammatory burden, and predicts the probability distribution of lesion grades across the rejection spectrum. These findings establish the three-gene signature as a noninvasive readout of biopsy-defined rejection lesions, demonstrate the analytical and interpretive advantages of first-order biomarker–lesion associations, and suggest a route toward integrating molecular urine testing with lesion-based risk stratification in kidney transplant care.

## RESULTS

### Study Cohort

Figure 1 summarizes the study schema and the analytic workflow. Among the 389 biopsy-urine pairs prospectively collected from kidney allograft biopsies, 35 (8.9%) were excluded because of BK virus nephropathy, pyelonephritis, or inadequate urinary RNA quantity or quality, yielding a final analytic cohort of 354 biopsy-urine pairs. All biopsies were processed according to standardized protocols and scored for Banff acute lesions by experienced renal pathologists. Total RNA was isolated from the corresponding urine sediment for absolute quantification of CD3E mRNA, CXCL10 mRNA, and 18S rRNA using customized RT-qPCR, and the CTOT-04 urinary cell three-gene diagnostic signature score was calculated for each sample using a validated equation ^19^.

**Figure 1.**
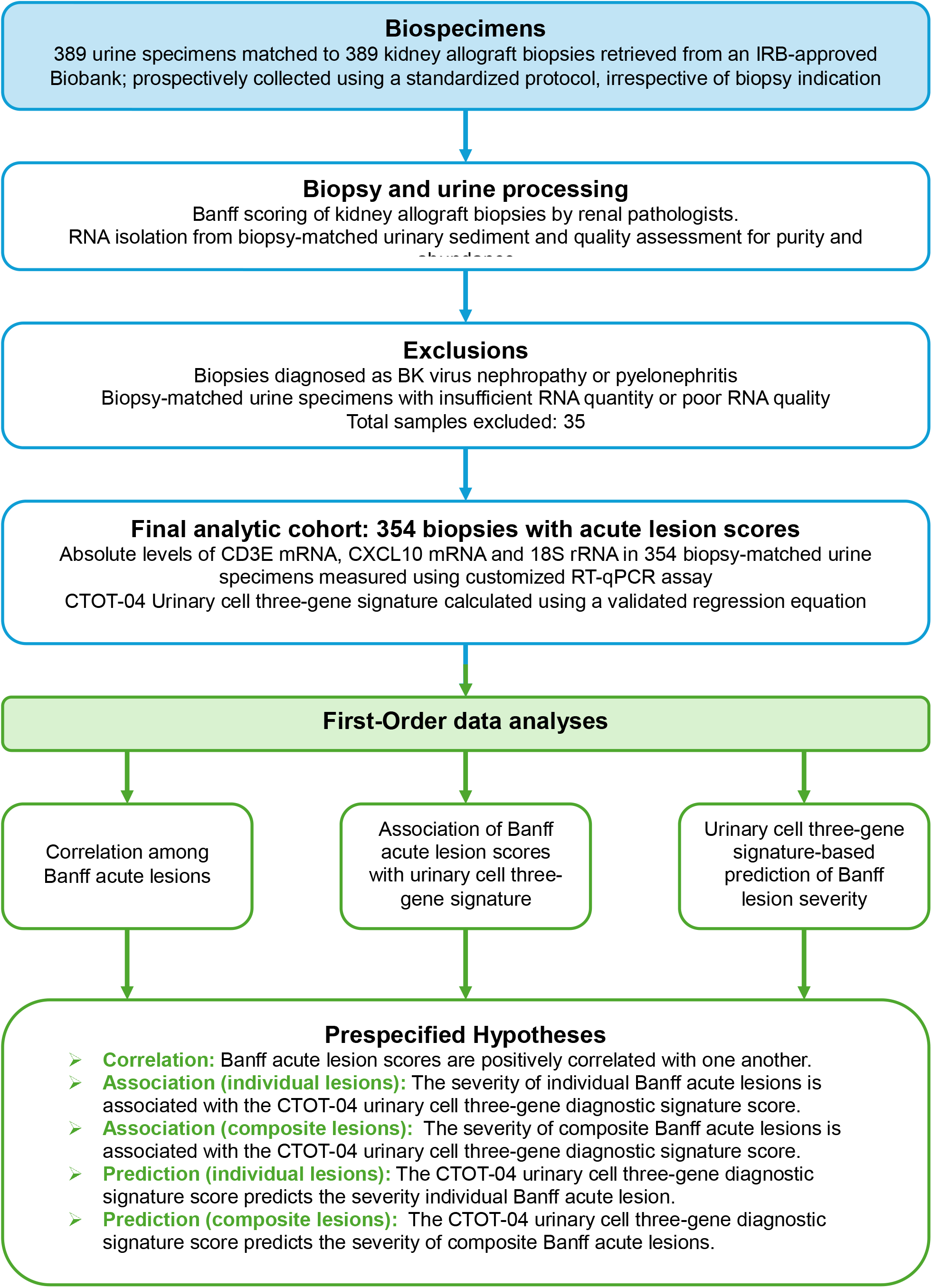
Study schema and analytic workflow. Flow diagram depicting the derivation of the final analytic cohort and the prespecified analyses. A total of 389 urine specimens matched to 389 kidney allograft biopsies, prospectively collected using a standardized collection protocol and stored in an IRB–approved biobank, were retrieved to determine the first-order association between Banff acute lesion scores and CTOT-04 urinary cell three-gene diagnostic signature score. The kidney allograft biopsies, performed for clinical indication or surveillance protocol biopsies, underwent Banff lesion scoring by experienced renal pathologists masked to the urinary cell three-gene score. Total RNA was isolated from biopsy matched urinary sediments and assessed for RNA quantity and quality. Biopsies diagnosed as BK virus nephropathy or pyelonephritis and urine specimens with insufficient or poor-quality RNA were excluded (total excluded, n=35), yielding a final analytic sample of 354 urine-biopsy with acute lesion scores. Absolute levels of CD3E mRNA, CXCL10 mRNA, and 18S rRNA were quantified in these 354 biopsy-matched urine specimens using a customized RT-qPCR assay, and the urinary cell 3-gene score was calculated using a previously validated regression equation (19). First-order analyses examined (i) correlations among Banff acute lesion scores, (ii) associations between individual and composite Banff acute lesion scores and the urinary cell three-gene score, and (iii) urinary cell three gene score–based prediction of Banff lesion severity. Prespecified hypotheses tested whether Banff acute lesion scores are positively correlated with one another, whether the severity of individual and composite Banff acute lesions is associated with the urinary cell three-gene score, and whether this three gene score predicts the severity of individual and composite Banff acute lesions.

Supplementary Table 1A-C summarizes the characteristics of the 302 kidney allograft recipients who contributed 354 biopsies, including 205 biopsies with Banff acute rejection (AR) and 149 without rejection (NR). Total RNA quantity and spectrophotometric purity indices were similar in AR and NR groups indicating that differences in urinary cell three-gene signature scores are unlikely to be driven by systematic variations in RNA recovery.

### Pairwise Associations Between Banff Lesion Scores

We first examined pairwise associations among *g, ptc, i,* and *t* using contingency tables with Goodman-Kruskal’s gamma and Somers’ D to test whether Banff acute lesion scores are positively correlated (Supplementary Table 2 and Supplementary Methods). The v lesion score was not considered as an independent category since v was scored as 0 in 95% of 354 biopsies, and biopsies with v lesions were already included in g, ptc, i, t categories.

The *g* and *ptc* scores were strongly associated (χ² 146.6; gamma 0.709; Somers’ D 0.567 for *ptc* given *g* and 0.416 for *g* given *ptc*), indicating that higher *g* scores occurred predominantly in biopsies with higher *ptc* score, consistent with their joint use to define microvascular inflammation. Associations of *g* with *i* and *t* were weaker but remained positive (gamma 0.340 and 0.325, respectively), suggesting that more severe glomerular inflammation tends to co-occur with higher tubulointerstitial scores, albeit with smaller effect size than for microvascular lesions. The *ptc* scores also showed moderate associations with *i* and *t* scores (gamma 0.556 and 0.590, respectively), indicating that peritubular capillaritis overlaps between both microvascular inflammation and tubulointerstitial inflammation.

The strongest relationship was between *i* and *t* (χ² 424.1; gamma 0.969; Somers’ D 0.702 for *t* given *i* and 0.874 for *i* given *t*), consistent with these two lesions reflecting a shared T-cell mediated inflammatory process, as formalized in the Banff criteria for T-cell mediated rejection.

Analyses of pair-wise associations; *g* by *ptc*, *g* by *i*, *g* by *t*, *ptc* by *i*, *ptc* by *t*, and *i* by *t,* demonstrate that Banff acute lesion scores are positively and strongly correlated with one another and form coherent microvascular inflammation and tubulointerstitial clusters (Supplementary Tables 3A-3F).

### Relationship of Individual Banff Lesion Scores with the CTOT-04 Urinary Cell Three-Gene Diagnostic Signature Score

We next examined the direct relationship between individual Banff acute lesion scores and the urinary three-gene score to test the hypothesis that the lesion severity is associated with the molecular signature.

Box-and-whisker plots (Figure 2) depict the urinary cell three-gene scores across the full range of ordinal lesion scores (0, 1, 2, and 3) for *g* (Figure 2A*), ptc* (Figure 2B)*, i* (Figure 2C) and *t* (Figure 2D). For each lesion score, a progressive increase in the three-gene score was evident with increasing histologic severity, indicating graded molecular activation that tracks lesion burden.

**Figure 2.**
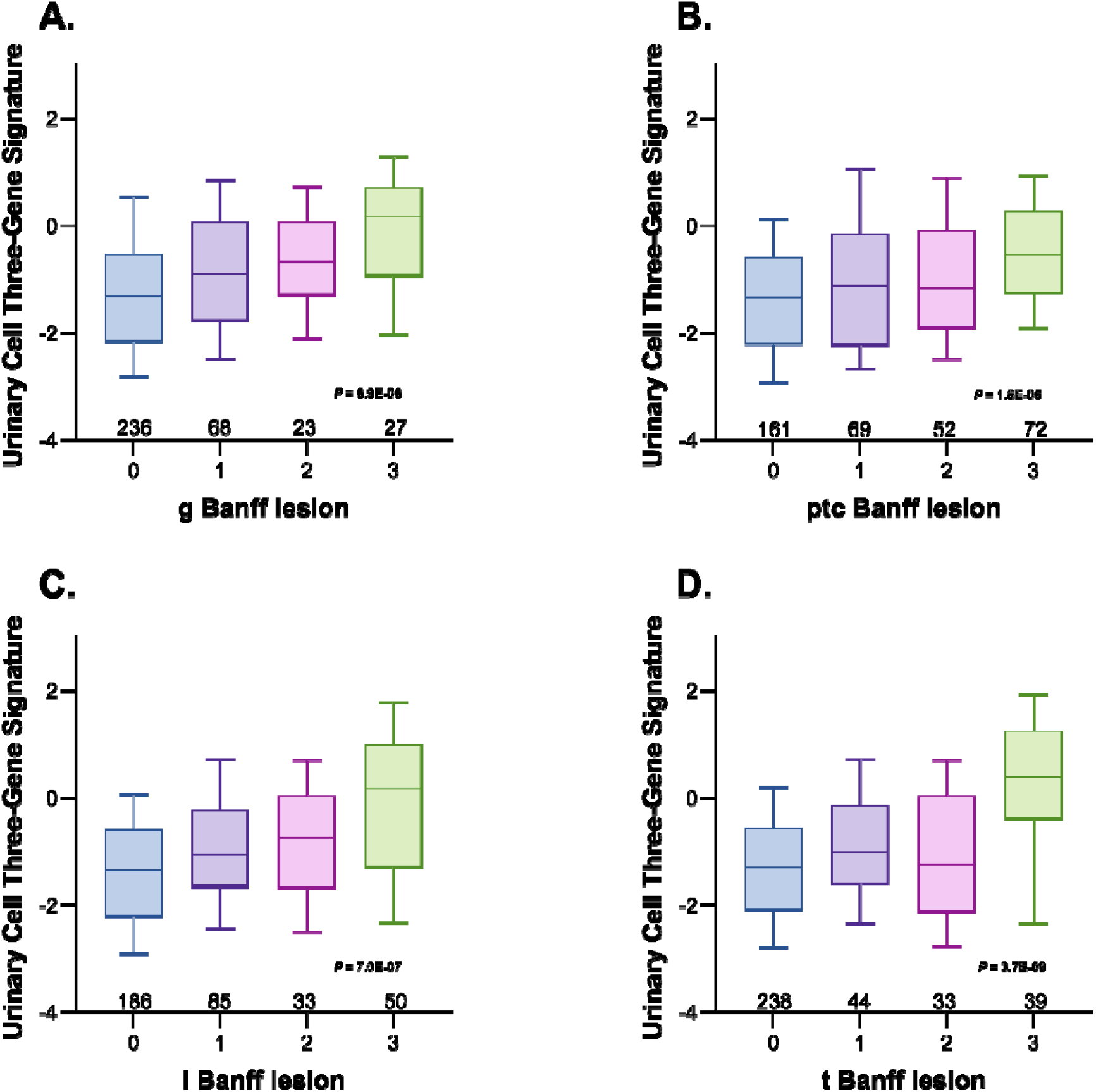
Box-and-whisker plots showing the distribution of the urinary cell three-gene signature scores for individual Banff acute lesion scores. The horizontal line within each box for glomerulitis (g, Panel A), peritubular capillaritis (ptc, Panel B), interstitial infiltration (i, Panel C), and tubulitis (t, Panel D), represents the median, the lower and upper borders of the box indicate the 25^th^ and 75^th^ percentiles, and the whiskers extend to inner fences, defined as 1.5 times the inter quarter range (IQR) below the 25^th^ percentile and above the 75^th^ percentile of the three-gene scores. The number (n) of biopsy-matched urine samples contributing to each Banff lesion score category is shown below the corresponding boxes. The urinary cell three-gene scores increased progressively with higher Banff scores for all four lesions, with significant global differences by Kruskal-Wallis test (g: P= 6.9E-06, ptc: P= 1.6E-05, i: P=7.0E-07, t: P=3.7E-09), greatest for tubulitis followed by and interstitial inflammation, glomerulitis and peritubular capillaritis. The effect size was largest for tubulitis followed by glomerulitis, interstitial inflammation, and peritubular capillaritis, after controlling for concurrent acute lesions (Table 1).

Global associations between each lesion score and the urinary cell three-gene score were evaluated using Kruskal-Wallis nonparametric test, which compares three-gene score distributions across ordered lesion categories without assuming normality, together with the _η_ -statistic from one-way ANOVA to quantify the proportion of variance in the three-gene score explained by lesion severity (Table 1). Detailed mean values and confidence intervals for three-gene scores across lesion categories are provided in Supplementary Table 4 and the rationale and strengths of this combined inferential and effect-size approach are provided in the Supplementary Methods.

**Table 1.**
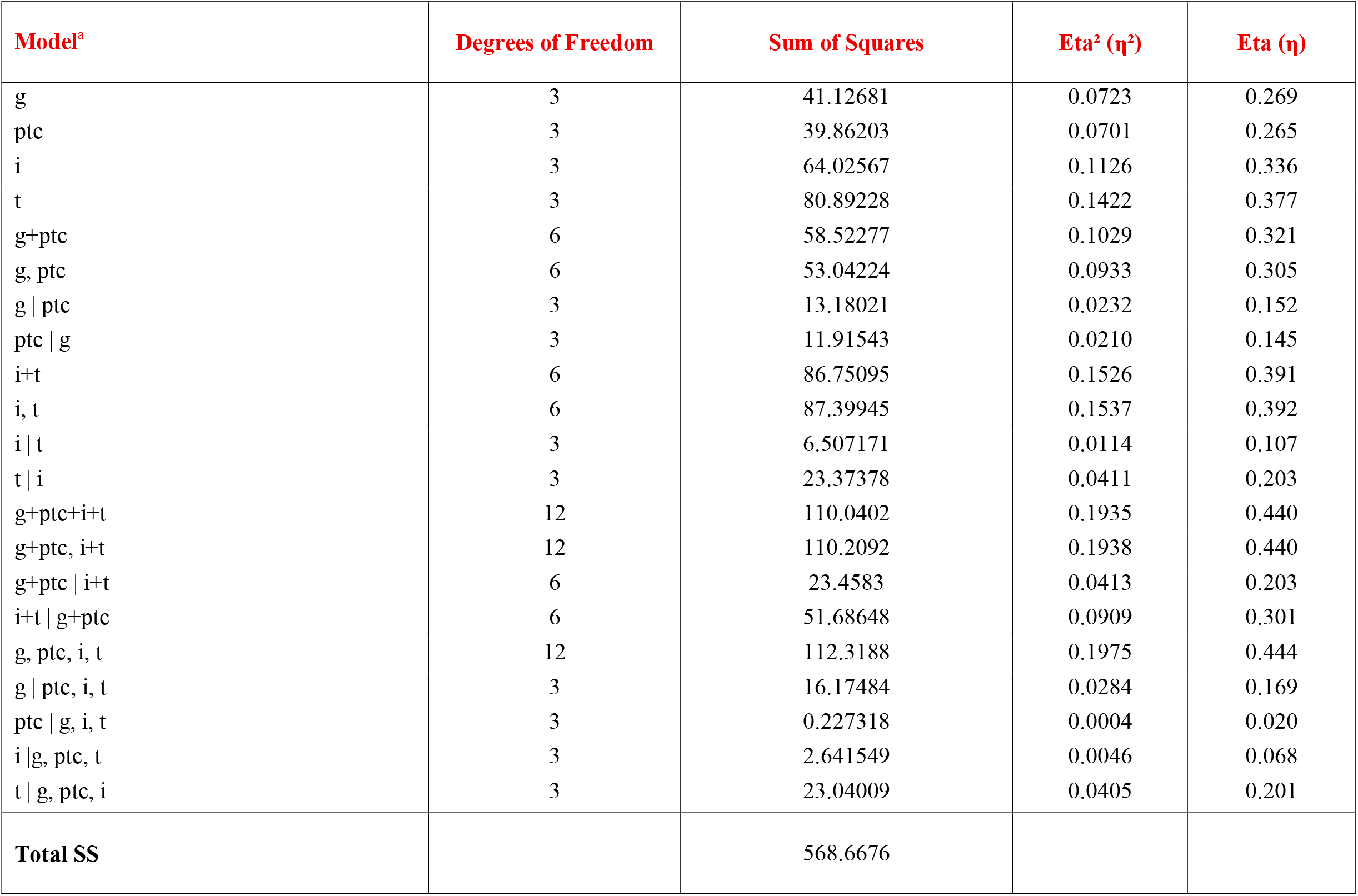
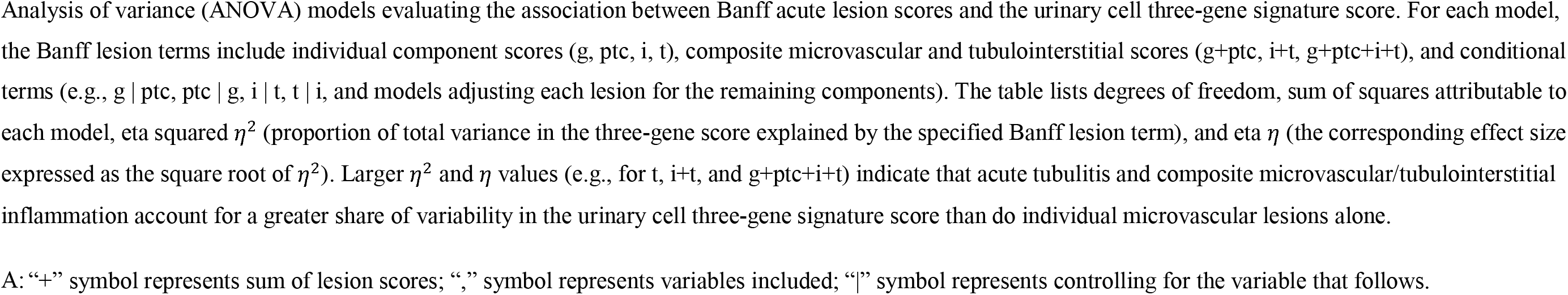
Analysis of Variance Models Predicting CTOT-04 Urinary Cell Three-Gene Diagnostic Signature Score from Banff Acute Lesion Scores.

For *g*, three-gene signature scores differed significantly across Banff g scores (P=6.9E-06, Figure 2), indicating that *g* severity accounts for meaningful fraction of the variance in the signature.

For *ptc*, the association was similar (P=1.6E-05, Figure 2), showing that differences in *ptc* severity explain a comparable proportion of variance in the gene scores.

Stronger associations were observed for the tubulointerstitial lesions: *i* was associated with significantly different urinary cell three-gene signature scores across lesion grades (P=7.0E-07, Figure 2), and *t* demonstrated the largest effect (P=3.7E-09, Figure 2), indicating that increasing *t* scores predict the greatest proportion of variance in the signature. While the urinary cell-three-gene signature score is strongly associated with all four Banff acute lesions, the effect size is greatest for tubulitis, followed by glomerulitis, interstitial inflammation, and peritubular capillaritis, after controlling for concurrent lesions (Table 1).

### General Linear Models Predicting the CTOT-04 Urinary Cell Three-Gene Signature Score from Individual Lesion Scores

To complement these nonparametric analyses and quantify the independent contribution of each lesion, we fitted general linear models (GLMs) with individual Banff acute lesion scores entered as categorical predictors and the urinary cell three gene score as the continuous outcome (Supplementary Table 4). Detailed rationale and strengths of this model-based approach are described in the Supplementary Methods.

In the model including *g* and *ptc* as indicators of microvascular inflammation, the combined lesion burden was significantly associated with the urinary cell three gene score (F □,□□□=5.95, P<0.001). Type III tests showed that, after adjusting for *ptc*, *g* remained significantly associated with the signature (F □,□□□=2.96, P=0.0325), indicating that mean three gene scores differ across *g* categories even after accounting for their correlation with ptc. Additionally, after adjusting for *g*, *ptc* remained significantly associated with the three gene score (F □,□□□=2.67, P=0.0473), indicating that *ptc* contributes independent information on the molecular signature beyond glomerular inflammation.

In a parallel GLM including *i* and tubulitis *t* as predictors, the urinary cell three-gene score was significantly associated with the combined tubulointerstitial lesion burden (F □,□□□=10.50, P<0.0001). After mutual adjustment, *t* remained strongly associated with the signature (Type III F □,□□□=5.62, P=0.0009), whereas *i* did not show an independent association (Type III F □,□□□=1.56, P=0.20). The mean three-gene scores were approximately 1–1.25 units higher in biopsies with t=3 compared with t=0–2 (all P≤0.014), indicating that the molecular signature increases markedly with severe tubulitis even after accounting for *i*.

### Relationship of Composite Banff Lesion Scores to CTOT-04 Urinary Cell Three-Gene Signature Score

We then investigated the association of composite Banff acute lesion scores that aggregate microvascular inflammation and tubulointerstitial inflammation, testing the hypothesis that the severity of composite indices is associated with the urinary cell three-gene score (Supplementary Table 5).

Global associations between composite lesion score and the urinary cell three-gene score were evaluated using Kruskal-Wallis nonparametric test. Detailed rationale and strengths of this combined inferential and effect-size approach are provided in the Supplementary Methods. Figure 3 and Table 1 show the first-order associations between the signature and composite scores representing microvascular inflammation (*g+ptc*) and tubulointerstitial inflammation (*i+t*). Figure 3A displays three-gene scores according to combined Banff *g+ptc* categories; Figure 3B displays gene signature scores according to the combined Banff *i+t* categories, reflecting intragraft tubuloinflammatory burden, and Figure 3C illustrates gene signature scores according to the combined *g+ptc+i+t* score, capturing both microvascular and tubulointerstitial inflammation burden.

**Figure 3.**
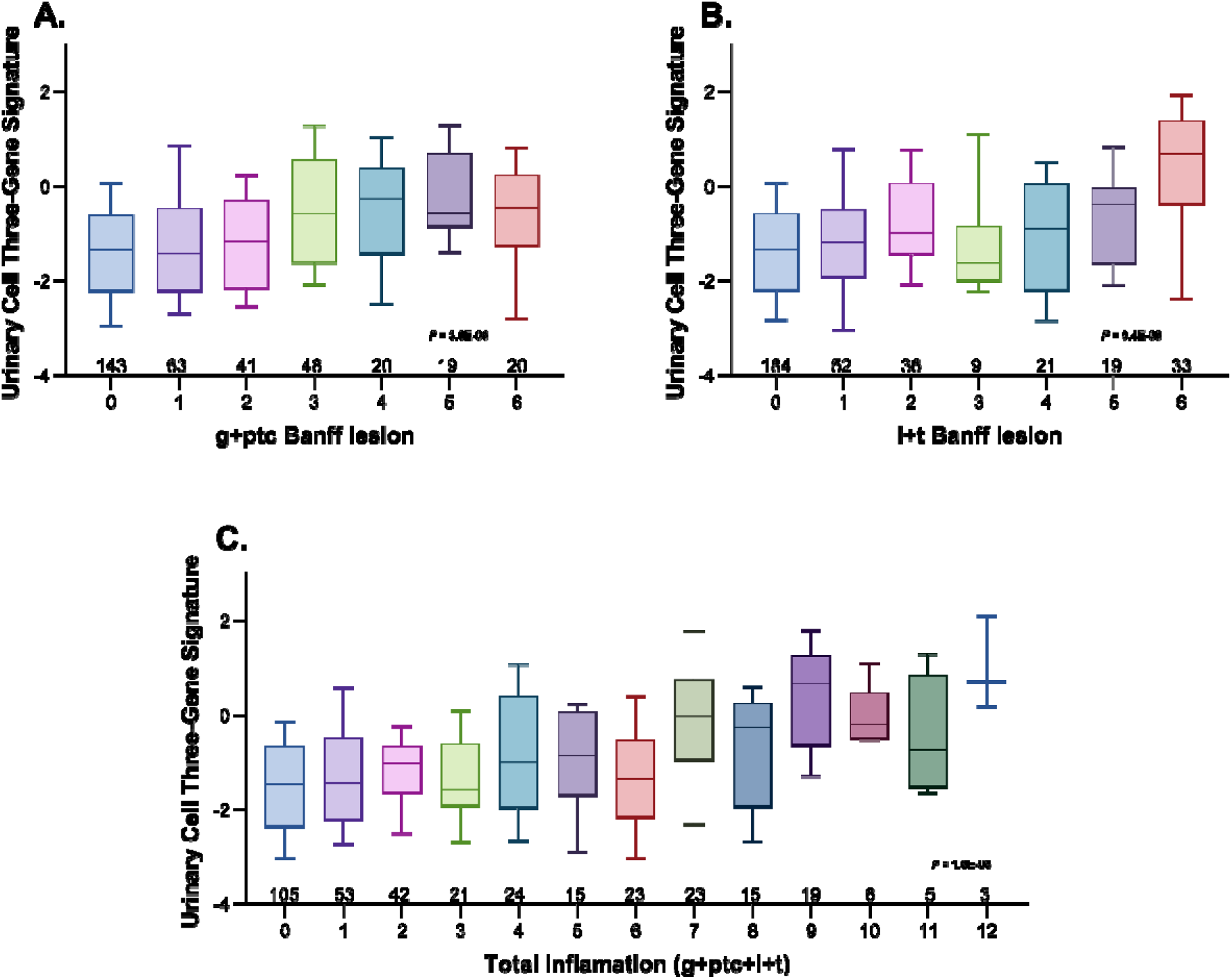
Box-and-whisker plots showing the distribution of the urinary cell three-gene signature scores for composite Banff acute lesion scores. The horizontal line within each box for glomerulitis plus peritubular capillaritis (g+ptc, Panel A), interstitial infiltration and tubulitis (i+t, Panel B) and g+ptc+i+t (Panel C), represents the median, the lower and upper borders of the box indicate the 25^th^ and 75^th^ percentiles, and the whiskers extend to inner fences, defined as 1.5 times the inter quarter range (IQR) below the 25^th^ percentile and above the 75^th^ percentile of the three-gene scores. The number of biopsy-matched urine samples contributing to each Banff lesion score category is shown below the corresponding boxes. Three-gene scores increased progressively with higher composite Banff scores for g+ptc (Panel A) i+t (Panel B), and g+ptc+i+t (Panel C) with significant global differences by Kruskal-Wallis test (g+ptc: P= 3.0E-06, Panel A), i+t P=9.4E-08, Panel B, g+ptc+i+t: P=1.6E-08, Panel C). The effect size was largest for g+ptc+i+t followed by tubulointerstitial inflammation (i,t) composite and (g+ptc) inflammation composite (Table 1).

Across composite histologic indices, higher three-gene scores were observed with increasing composite severity, indicating a graded relationship between the biopsy-based inflammatory activity and urinary cell three-gene signature. Associations were statistically significant by Kruskal-Walis one-way analyses: *g+ptc* (P=3.0E-06, Figure 3A), *i+t* (P=9.4E-08, Figure 3B), and *g+ptc+i+t* (P=1.6E-08, Figure 3C). These data reported in Table 1 support that the substantial variation in both microvascular and tubulointerstitial inflammation is captured by the urinary cell three-gene signature, and that aggregate inflammatory burden is particularly well reflected in the molecular readout.

The magnitude of association of urinary cell three-gene signature was greater for the *i+t* composite than the *g+ptc* composite, as reflected by both the smaller P value (9.4E-08 versus 3.0E-06) and higher Eta value for *i+t* (0.391 versus 0.321). This pattern suggests that the signature may align more strongly with tubulointerstitial inflammatory lesions than with the isolated microvascular inflammatory lesions, although both relationships were robust and biologically meaningful.

### General Linear Model Predicting the CTOT-04 Urinary Cell Three-Gene Signature Score from Composite Lesion Scores

To assess whether the composite microvascular and tubulointerstitial indices contribute independently to the urinary cell three-gene signature, we fitted a GLM with the urinary cell three-gene score as the dependent variable and both *g+ptc* and *i+t* composite scores as predictors. GLM results from the composite lesion models are summarized in Supplementary Table 5.

Controlling for *i+t*, the *g+ptc* composite remained significantly associated with the three-gene score (F_6,341_=2.91, P=0.0089), indicating that the mean urinary cell three-gene values differ across *g+ptc* categories even after accounting for their association with tubulointerstitial inflammation.

Controlling for *g+ptc*, the *i+t* composite accounted for a larger proportion of the variance in the three-gene score (F_6,341_=6.41, P<0.0001), demonstrating a particularly strong graded relationship between the gene signature and parenchymal cellular inflammation. Together with the Kruskal-Wallis results, these findings support a robust, model-based association between increasing severity of both microvascular (*g+ptc*) and tubulointerstitial (*i+t*) inflammatory injury and higher urinary cell three-gene signature scores.

Finally, we examined the urinary cell three-gene signature as a function of all four Banff lesion scores (*g, ptc, i,* and *t*) entered simultaneously as four-level categorical predictors (0–3). The overall model was statistically significant (F□,□□□=6.99, P<0.0001) and explained about 20% of the variability in the three-gene score, indicating that the combined histologic lesions account for a meaningful fraction of the molecular signal. After mutual adjustment, the Type III tests showed that *g* and *t* remained independently associated with the signature score (*g*: F □,□□□=4.03, P=0.0078; *t*: F □,□□□=5.74, P=0.0008), whereas *ptc* and *i* did not contribute additional explanatory power (*ptc*: F □,□□□=0.06, P=0.98; *i*: F □,□□□=0.66, P=0.58). Consistent with this, mean three-gene scores were significantly higher in biopsies with *g*=3 or *t*=3 compared with lower grades: relative to *g*=3, categories *g*=0 and *g*=1 had three-gene scores lower by 0.90 and 0.63 units, respectively (P≤0.03), and relative to *t*=3, categories *t*=0–2 had scores lower by approximately 1.0–1.25 units (P≤0.015). These findings indicate that, when all acute Banff lesions are considered together, the signature is most strongly and independently linked to severe tubulitis and glomerulitis.

### CTOT-04 Urinary Cell Three-Gene Diagnostic Signature Score Predicting Banff Lesion Severity

Given the strong associations between Banff acute lesion scores and the urinary cell three-gene signature, we next evaluated this molecular score as a noninvasive predictor of histologic lesion severity. The rationale and strengths of the predictive modeling framework are detailed in the Supplementary Methods.

In separate proportional-odds ordinal logistic regression models, each Banff acute lesion score (0,1, 2 or 3) was modeled as an ordinal outcome with the three-gene signature core as the primary predictor. Plots of the three-gene score versus predicted lesion probability (Figure 4A–D) show a graded increase in the probability of having ≥1, ≥2, or 3 lesion score for *g, ptc, i*, and *t* as the three-gene score increases, consistent with a strong monotonic relationship between the molecular signature and acute injury burden.

**Figure 4.**
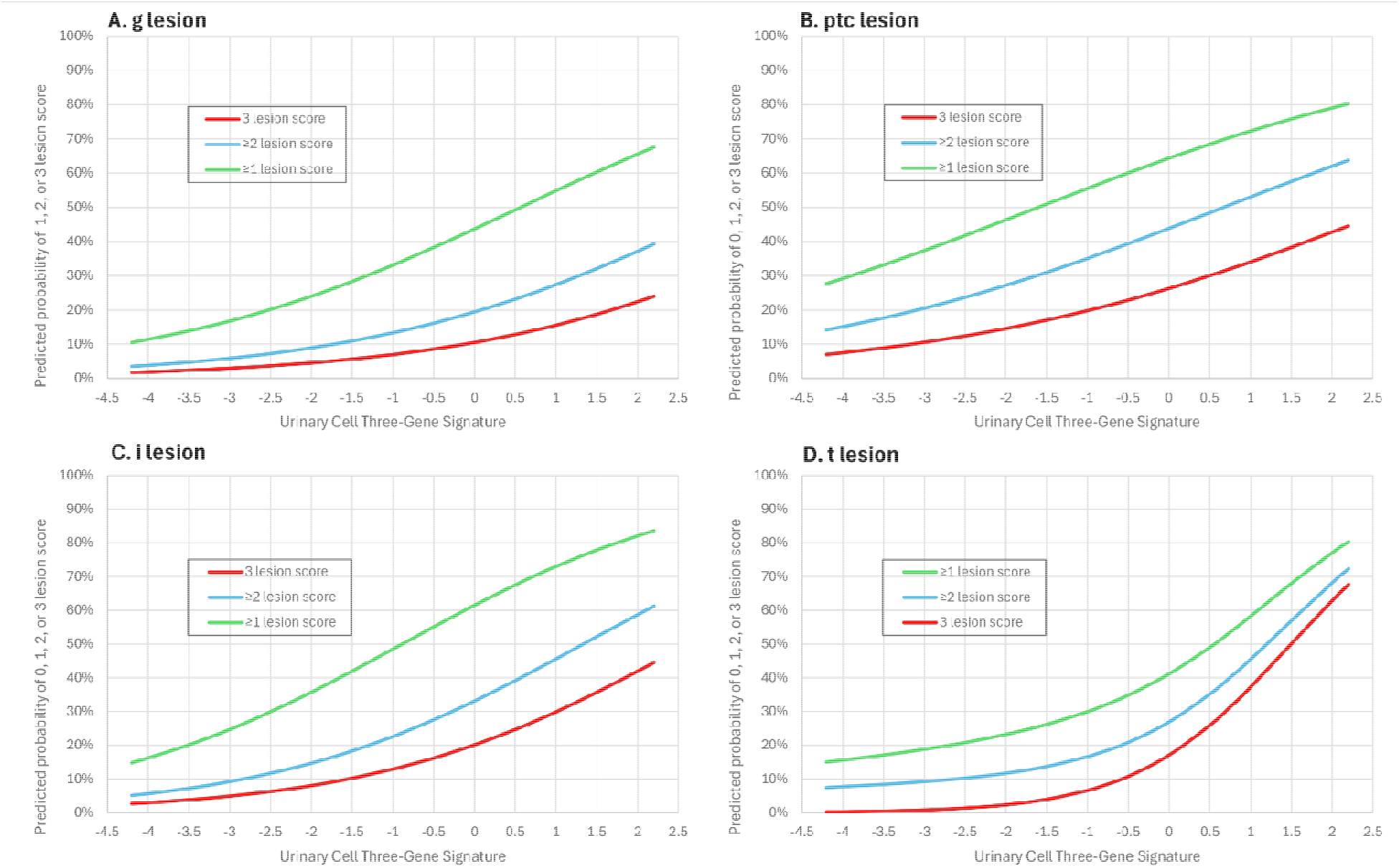
Association of CTOT-04 urinary cell three-gene diagnostic signature score with the severity of Banff acute lesions in kidney allograft biopsies. Predicted probabilities from proportional-odds ordinal logistic regression models relate the continuous urinary cell three-gene score to the ordinal severity scores for Banff glomerulitis (g, Panel A), peritubular capillaritis (ptc, Panel B), and interstitial inflammation (i, Panel C). Because tubulitis (t, Panel D) violated the proportional-odds assumption (Table 2A), the predicted probabilities for t were obtained from a non-ordinal generalized logistic regression model (Table 2B). For each lesion, curves show the model-based probability of having at least a mild lesion (grade ≥1, green), at least a moderate lesion (grade ≥2, blue), and a severe lesion (grade 3, red) across the observed range of urinary cell three-gene scores. The plots showing increasing urinary cell three-gene scores are associated with progressively higher probabilities of grade ≥1, ≥2, and 3 lesions, indicate that the urinary cell gene-expression signature closely tracks the severity of Banff acute lesions in kidney allograft biopsies.

Regression analyses, summarized in Table 2, quantify these associations. In the single-lesion models, each 1-unit increase in the signature was associated with significantly higher odds of more severe lesions, with odds ratios (95% CI) of 1.57 (1.31–1.88) for *g*, 1.45 (1.23–1.69) for *ptc*, 1.70 (1.43–2.01) for *i,* and 1.79 (1.48–2.16) for *t* (all P<0.0001) (Table 2A). Composite lesion scores showed similarly elevated odd ratios per unit increase in signature: 1.55 (1.32–1.81) for *g+ptc,* 1.71 (1.45–2.02) for *i+t*, 1.76 (1.51-2.06) for *g+ptc+i+t* (all P<0.0001). Likelihood-ratio χ² statistics for the gene score (df=1) were 24.41, 22.85, 38.31, 37.22, 31.92. 39.86 and 52.00 for *g*, *ptc, i, t, g+ptc, i+t, g+ptc+i+t*, respectively (all P<0.0001), indicating that the signature adds substantial discriminatory information across categories of both microvascular and tubulointerstitial injury.

**Table 2.** Regression Models Predicting Banff Acute Lesion Score From CTOT-04 Urinary Cell Three-Gene Diagnostic Signature Scores.

Overall calibration of these ordinal models, assessed by the Hosmer–Lemeshow goodness-of-fit test (Table 2A), was acceptable, with χ² (df=26) and P values of 20.71 and 0.76 for *g*, 31.78 and 0.20 for *ptc*, 28.76 and 0.32 for *i,* and 34.42 and 0.12 for *t*. For composite scores, χ² (df=53) of 49.48 (P=0.61) for *g+ptc* and 76.08 (P=0.0205) for *i+t,* and χ² (df=107) of 108.34 (P=0.44) for *g+ptc+i+t*, indicating good model fit for all outcomes except *i+t* where modest miscalibration was observed.

**Table 2A.**
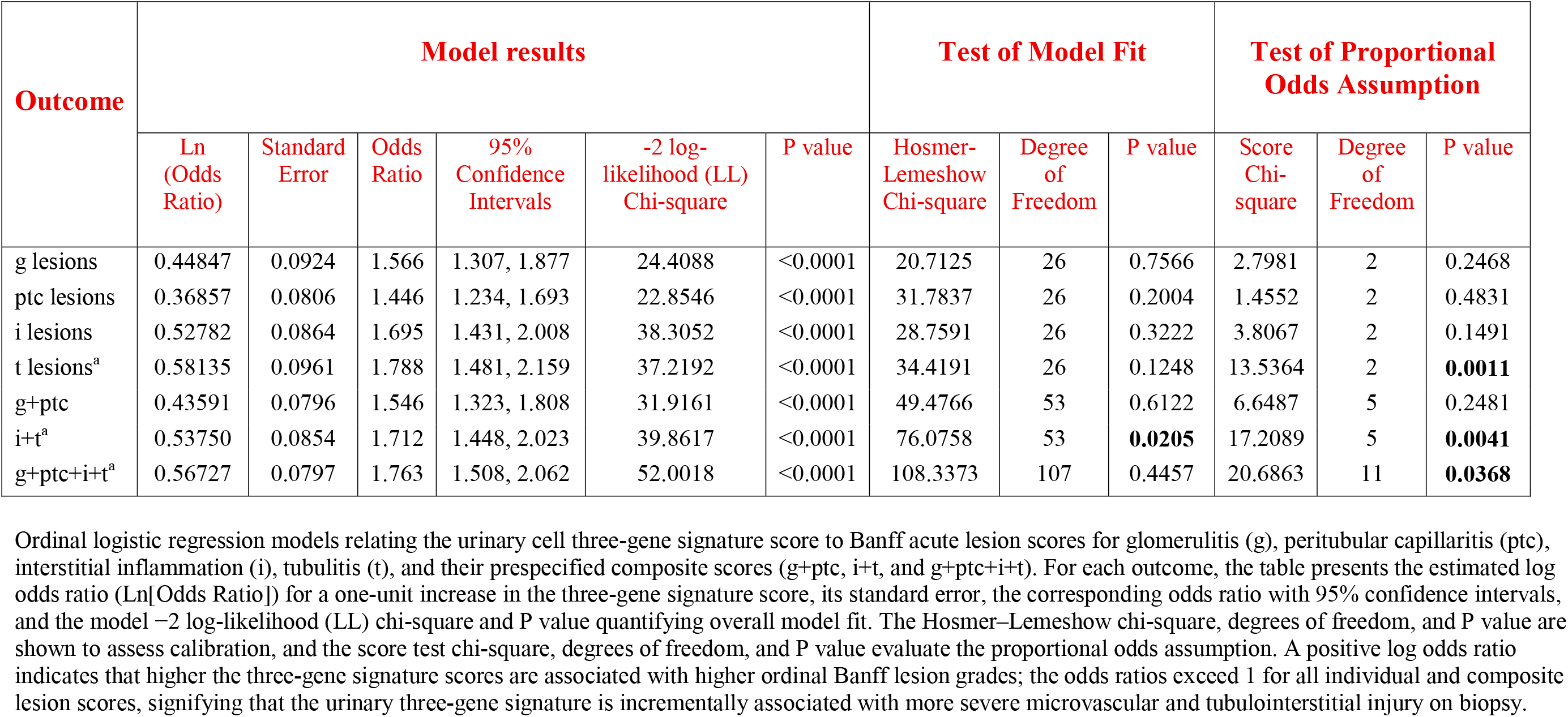
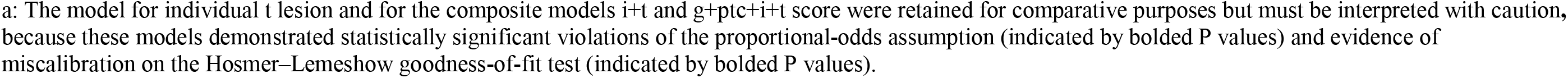
Ordinal Logistic Regression Models.

| Outcome | Model results |  |  |  |  |  | Test of Model Fit |  |  | Test of Proportional Odds Assumption |  |  |
| --- | --- | --- | --- | --- | --- | --- | --- | --- | --- | --- | --- | --- |
|  | Ln (Odds Ratio) | Standard Error | Odds Ratio | 95% Confidence Intervals | -2 log-likelihood (LL) Chi-square | P value | Hosmer-Lemeshow Chi-square | Degree of Freedom | P value | Score Chi-square | Degree of Freedom | P value |
| g lesions | 0.44847 | 0.0924 | 1.566 | 1.307, 1.877 | 24.4088 | <0.0001 | 20.7125 | 26 | 0.7566 | 2.7981 | 2 | 0.2468 |
| ptc lesions | 0.36857 | 0.0806 | 1.446 | 1.234, 1.693 | 22.8546 | <0.0001 | 31.7837 | 26 | 0.2004 | 1.4552 | 2 | 0.4831 |
| i lesions | 0.52782 | 0.0864 | 1.695 | 1.431, 2.008 | 38.3052 | <0.0001 | 28.7591 | 26 | 0.3222 | 3.8067 | 2 | 0.1491 |
| t lesions<sup>a</sup> | 0.58135 | 0.0961 | 1.788 | 1.481, 2.159 | 37.2192 | <0.0001 | 34.4191 | 26 | 0.1248 | 13.5364 | 2 | **0.0011** |
| g+ptc | 0.43591 | 0.0796 | 1.546 | 1.323, 1.808 | 31.9161 | <0.0001 | 49.4766 | 53 | 0.6122 | 6.6487 | 5 | 0.2481 |
| i+t<sup>a</sup> | 0.53750 | 0.0854 | 1.712 | 1.448, 2.023 | 39.8617 | <0.0001 | 76.0758 | 53 | **0.0205** | 17.2089 | 5 | **0.0041** |
| g+ptc+i+t<sup>a</sup> | 0.56727 | 0.0797 | 1.763 | 1.508, 2.062 | 52.0018 | <0.0001 | 108.3373 | 107 | 0.4457 | 20.6863 | 11 | **0.0368** |

Ordinal logistic regression models relating the urinary cell three-gene signature score to Banff acute lesion scores for glomerulitis (g), peritubular capillaritis (ptc), interstitial inflammation (i), tubulitis (t), and their prespecified composite scores (g+ptc, i+t, and g+ptc+i+t). For each outcome, the table presents the estimated log odds ratio (Ln[Odds Ratio]) for a one-unit increase in the three-gene signature score, its standard error, the corresponding odds ratio with 95% confidence intervals, and the model -2 log-likelihood (LL) chi-square and P value quantifying overall model fit. The Hosmer-Lemeshow chi-square, degrees of freedom, and P value are shown to assess calibration, and the score test chi-square, degrees of freedom, and P value evaluate the proportional odds assumption. A positive log odds ratio indicates that higher the three-gene signature scores are associated with higher ordinal Banff lesion grades; the odds ratios exceed 1 for all individual and composite lesion scores, signifying that the urinary three-gene signature is incrementally associated with more severe microvascular and tubulointerstitial injury on biopsy.
a: The model for individual t lesion and for the composite models i+t and g+ptc+i+t score were retained for comparative purposes but must be interpreted with caution, because these models demonstrated statistically significant violations of the proportional-odds assumption (indicated by bolded P values) and evidence of miscalibration on the Hosmer–Lemeshow goodness-of-fit test (indicated by bolded P values).

The proportional odds (parallel slopes) assumption, evaluated by the score test, held for *g*, *ptc*, *i*, and *g+ptc* (χ²[df=2–5] 1.46–6.65, all P≥0.15), but was rejected for *t* (χ²=13.54, df=2, P=0.001), *i+t* (χ²=17.21, df=5, P=0.004), and *g+ptc+i+t* (χ²=20.69, df=11, P=0.037), indicating that a single pooled odds ratio does not fully capture the relationship between the three-gene score and tubulitis or composite lesion severity that includes *t* lesion (Table 2A). Accordingly, for the *t* lesion, we additionally fitted a generalized (multinomial) logistic regression model to obtain category-specific effects of the signature across Banff *t* grades 0–3 (Table 2B). This model showed marked declines in the predicted probability of *t*=0 and steep increases in the probability of *t*=3 with higher three-gene scores, reinforcing the concept that this noninvasive signature closely tracks key acute rejection lesions. In line with these analyses, Figure 4D is based on plots on generalized logistic regression, whereas Figure 4A-C are based on ordinal logistic regression models.

**Table 2B:** Generalized Logistic Regression Analysis.

| Outcome | Predictor |
| --- | --- |
| Banff t Lesion Score, t0 as reference | Three-Gene Score, OR (95% CI) |
| t3 vs. t0 | 3.082 (2.174, 4.367) |
| t2 vs. t0 | 1.174 (0.863, 1.597) |
| t1 vs. t0 | 1.269 (0.966, 1.667) |
| -2 LL chi-square, df=3 | 53.6265 |
| P value | <0.0001 |
| Hosmer-Lemeshow goodness-of-fit test, chi-square, df=26 | 30.1803 |
| P value | 0.1788 |

Generalized logistic regression model with t0 as reference; odds ratios (OR) and 95% confidence intervals (CI) are shown for the effect of a one-unit increase in the urinary cell three-gene score on the odds of being in each t category (t1, t2, t3) versus t0. The $-2 \log \text{likelihood}$ ( $-2 \text{ LL}$ ) $\chi^2$ (df=3) value 53.6265 and its P value <0.0001 show the significant contribution of the urinary cell three-gene score to the model. Collectively, the pattern shows a monotonic increase in effect size with higher t lesion grade, with the strongest and clearly significant association for severe t3 tubulitis. The Hosmer–Lemeshow goodness-of-fit $\chi^2$ (df=26) value 30.1803 and its P value 0.1788 show the adequacy of model fit and suggest that the generalized logistic regression model adequately describes the relationship between the urinary cell three-gene score and the distribution of Banff t lesion scores.

## DISCUSSION

In this single-center study of 354 biopsy-urine pairs, the CTOT-04 urinary cell three-gene diagnostic signature emerged as a quantitative, noninvasive readout of Banff acute lesion biology. The score increased progressively across Banff 0–3 *for g, ptc, i* and *t*, and general linear, ordinal and generalized logistic models showed that higher signature values were associated with substantially increased odds of more severe histologic injury, even after adjustment for concurrent lesions. These results demonstrate that the urinary cell three-gene score behaves as a continuous molecular correlate of intragraft inflammatory burden and directly encodes the severity of rejection related tissue injury.

A central conceptual advance of this work is the explicit focus on first-order associations between a urinary cell biomarker and individual Banff lesion scores, rather than second order associations with composite diagnostic categories such as TCMR or ABMR. Banff diagnostic categories integrate multiple lesion scores alongside clinical and serologic features and are therefore heterogeneous and potentially confounded; hence, biomarker–diagnosis correlations may obscure the specific histologic processes that the biomarker measures. By anchoring our analyses directly to *g, ptc, i* and *t*, the present study links the urinary cell three-gene signature to the quantitative histologic features that drive diagnostic assignment and aligns transplant biomarker evaluation with contemporary frameworks that prioritize mechanistic interpretability and analytic transparency ^24,25^.

The pattern of associations observed is biologically coherent and extends prior work on urinary cell transcriptomics ^31^. The CD3E–CXCL10–18S signature reflects intragraft T-cell activation and interferon-γ–inducible chemokine signaling; in this cohort, the same locked three gene score tracked both tubulointerstitial injury (*i* and *t*) and microvascular inflammation (*g* and *ptc*), with particularly strong relationships *for i, t* and the composite indices *i+t* and *g+ptc+i+t*. These findings are consistent with accumulating evidence that T cell–mediated and antibody mediated rejection share overlapping inflammatory pathways, including T cell infiltration and chemokine driven recruitment of effector cells into both interstitial and microvascular compartments ^31–33^. The robust associations with *g+ptc*, i*+t* and *g+ptc+i+t* indicate that the urinary cell signature reflects global rejection related inflammatory activity across compartments rather than a single diagnostic category and therefore has the potential to serve as a integrative molecular readout for heterogeneous rejection phenotypes.

The modeling strategy used here clarifies how the urinary cell three-gene score relates to lesion severity and distribution. Nonparametric analyses documented monotonic increases in scores across lesion categories, establishing a graded molecular response with rising histologic injury. General linear models and corresponding analysis of variance quantified the proportion of variance in the molecular signal explained by microvascular versus tubulointerstitial inflammation and showed that tubulitis and composite microvascular/tubulointerstitial indices account for the largest share of variability. Ordinal and generalized logistic regression then translated these associations into clinically interpretable effect estimates: each 1-unit increase in the signature was associated with roughly 40–70% higher odds of being in a more severe lesion category, with the strongest effects for *t* and for the *i+t* and *g+ptc+i+t* composites. Taken together, these complementary analyses show that the urinary three-gene score provides independent information beyond any single Banff lesion and functions as a continuous molecular summary of histologic rejection burden.

This study also illustrates the feasibility and value of a locked-equation approach for urinary cell RT-qPCR biomarkers. The CTOT-04 urinary cell three gene diagnostic signature score was calculated using coefficients from the original trial without recalibration in this cohort, mirroring real-world implementation and minimizing the risk of overfitting ^19^. The strong, graded first order associations across diverse lesion patterns suggest that the biology captured by CD3E, CXCL10 and 18S is conserved across case mixes and that this signature can be deployed as a standardized quantitative readout in multicenter studies and clinical trials.

Several limitations warrant consideration. Ours is a single center study, and external validation is essential to establish generalizability across laboratories, platforms and pathologists. Although biopsies with BK polyomavirus nephropathy or inadequate RNA were excluded, other forms of graft injury—including calcineurin inhibitor toxicity, chronic scarring and recurrent native kidney disease—may have influenced both histology and urinary transcripts. The cross-sectional design focuses on contemporaneous biopsy–urine pairs and does not capture temporal dynamics, although prior work suggests that the CTOT-04 urinary cell three gene diagnostic signature score may rise before histologic rejection becomes apparent ^19^. Finally, Banff lesion scoring, even when performed by experienced transplant pathologists, is subject to inter-observer variability ^34,35^, which would be expected to bias biomarker–lesion associations toward the null rather than inflate them.

Despite these limitations, the findings have several implications for biomarker development, clinical trial design and future practice. First, they provide mechanistic support for using the urinary cell three gene signature as an intermediate endpoint or surrogate biomarker in studies targeting intragraft inflammation, particularly in settings where protocol biopsies are impractical or carry unacceptable risk. Second, the strong relationships with both microvascular and tubulointerstitial composites indicate that the signature could be incorporated into multivariable risk models that integrate molecular, serologic and histologic data to refine rejection phenotyping, prognosis and treatment decision-making. Third, the first order analytic framework introduced here offers a generalizable template for evaluating emerging biomarkers by grounding validation in quantitative lesion scores rather than exclusively in composite diagnostic categories, thereby improving interpretability and alignment with the biology of tissue injury.

Activity and chronicity indices derived from Banff lesion scores are increasingly emphasized in recent Banff work plans and reports, reflecting a broader shift toward continuous, lesion based measures of rejection burden ^14,15^. Studies of automated histologic classification ^13^, Banff based chronicity indices in antibody mediated rejection ^11,12^ and large multicenter biopsy cohorts ^14^ have shown that lesion based indices more accurately capture the rejection spectrum and more precisely predict graft failure than conventional diagnostic labels. The present results extend this evolving paradigm by demonstrating that a noninvasive urinary molecular signature is tightly linked to these lesion-level metrics, effectively bridging histologic and molecular assessments of rejection activity.

In conclusion, the CTOT-04 urinary cell three gene diagnostic signature shows robust, graded first-order associations with individual and composite Banff acute lesion scores in kidney allograft biopsies. These data establish that the noninvasive molecular signal closely tracks the histologic substrates of rejection, support its biological plausibility as a surrogate of intragraft inflammation and highlight the methodological and translational advantages of focusing on direct biomarker–lesion associations in transplant biomarker research with immediate clinical applicability.

## METHODS

### Study Cohort

We analyzed 354 kidney allograft biopsy specimens with matched urine samples collected prospectively from 302 kidney transplant recipients. The study cohort was enrolled under an Institutional Review Board (IRB) -approved protocol. All participants provided written informed consent and underwent biopsies at our hospital, NewYork Presbyterian/Weill Cornell Medicine, and received follow-up standardized post-transplant care at our hospital.

Biopsies were performed either for clinical indication (elevated serum creatinine, proteinuria or other clinical concerns) or as part of protocol surveillance. Urine samples were collected within 24 hours of biopsies in all but 2 cases to ensure temporal correlation between urinary biomarker measurement and histopathological findings. Exclusion criteria included urinary tract infection at the time of sample collection, BK polyomavirus nephropathy, inadequate biopsy tissue for Banff scoring and insufficient urinary cell pellet for RNA extraction (N=35, not included in N=354).

Clinical variables were abstracted from a prospectively maintained, IRB-approved research database and cross-checked against electronic medical records for accuracy. The study was conducted in accordance with the ethical principles of the Declaration of Helsinki.

### Banff Pathology Assessment

Percutaneous needle core biopsies were obtained under ultrasound guidance. All kidney allograft biopsies were processed according to standardized protocols and evaluated by transplant pathologists using Banff classification criteria. Banff acute lesions were scored (0-3) for the following: *g*, *ptc*, ***i***, and *t*. The v lesion score was not considered as an independent category since v was scored as 0 in 95% of 354 biopsies, and biopsies with v lesions were already included in g, ptc, i, t categories. Pathologists were masked to CTOT-04 signature score. Cross-tabulation was used to quantify the strength and direction of associations among the Banff lesion scores.

### Urinary Cell mRNA Quantification by Customized RT-qPCR

Approximately 50 ml of urine and was processed at our Gene Expression Monitoring (GEM) Core. Of the 354 biopsy-matched urine samples, 330 were collected on the day of biopsy, 22 within one day after the procedure, and 2 within 3 days after biopsy.

Urine was centrifuged at 2000g for 30 min at room temperature to generate urinary cell pellets. Sedimented cells were lysed in a mixture of RNAlater (50 μl), RLT buffer and Beta-mercaptoethanol (350 μl), and total RNA was extracted using the RNeasy mini kit (Qiagen, Cat. 74,104). RNA purity (A260/A280) and yield (A260) were determined using a NanoDrop ND-1000 spectrophotometer (ThermoFisher Scientific).

Total RNA was reverse transcribed (RT) to cDNA with the TaqMan reverse transcription kit (Applied Biosystems Cat. N808–0234), as described previously (19) and as detailed in Supplemental Methods.

Gene-specific oligonucleotide primers and TaqMan fluorogenic (hydrolysis) probes, designed in our laboratory, were used to quantify mRNA for CD3E, CXCL10, TGF-B1 and 18S rRNA.

To address the low RNA yield from urinary cells, and enable measurement of multiple mRNAs, we employed a preamplification protocol developed in our laboratory, as described previously and as detailed in Supplemental Methods. Absolute transcripts copy number was determined using a universal standard curve generated from an in-house synthesized Bak amplicon. The three -gene signature score was calculated using a locked regression equation derived from our previously published CTOT-04 study: Three-gene score = -6.1487+ 0.8534 log_10_ (CD3E /18S) + 0.6376 log_10_ (CXCL10/18S) + 1.6464 log_10_(18S).

#### Statistical Analysis

Continuous variables are presented as mean (SD) and median (interquartile range) and categorical variables as frequencies and percentages. The distribution of the three-gene signature scores across Banff lesion scores was visualized using box-and-whisker plots.

Each Banff acute lesion score was modeled as an ordinal outcome using proportional-odds logistic regression with CTOT-04 urinary cell three-gene signature score as the primary predictor. To quantify the incremental contribution of the gene signature beyond a baseline model, we compared nested models using likelihood-ratio tests based on changes in the log-likelihood. The proportional odds (parallel slopes) assumption was evaluated for each lesion using a score test; when this assumption did not hold for a given lesion, we fitted a generalized logistic regression model that relaxed the proportionality odds constraint for the three-gene signature. The rationale and strengths of the predictive modeling framework are detailed in the Supplementary Methods.

## DATA AVAILABILITY

The source data is provided with this manuscript.

## ACKNOWLEDGEMENTS

We are grateful to our patients and our clinical colleagues. We thank our research coordinators Ms. Isha Bal and Ms. Mansi Milan Mhatre for their expert help.

## AUTHOR CONTRIBUTIONS

**MS** designed research studies

**CL** conducted experiments

**CL, TS, DMD, ADV, WH, SS, SVS, VKS, TM** acquired data

**CL, JES, TM** analyzed data

**MS** provided reagents

**JES, TM, MS** wrote the manuscript

All the authors participated in data interpretation and critically reviewed the manuscript.

