## Supplementary Materials for "First Order Associations Between Banff Acute Lesions in Kidney Allograft Biopsies and a Urinary Cell Three-Gene Diagnostic Signature"

**SUPPLEMENTARY INFORMATION**

**Table of Contents**

**Pages**

**Supplementary Methods**-----------------------------------------------------------------------------**3 - 8**

Study Cohort-------------------------------------------------------------------------------------------3

Banff Pathology Assessment of Kidney Allograft Biopsies-------------------------------------3

Urinary Cell mRNA Quantification BY Customized RT-qPCR--------------------------------3

Rationale and strengths of statistical testing-------------------------------------------------------5

**Supplementary References**--------------------------------------------------------------------------**9**

**Supplementary Tables**-------------------------------------------------------------------------------**10 - 34**

**Supplementary Table 1A.** Demographics and Clinical Characteristics of

Adult Kidney Allograft Recipients----------------------------------------------------------------	10 - 11

**Supplementary Table 1B.** Allograft Biopsy Histopathology Characteristics--------------	12

**Supplementary Table 1C.** Urinary Cell RNA Characteristics-------------------------------	13

**Supplementary Table 2.** Summary Statistics Crosstabulations for

each pair of lesion scores---------------------------------------------------------------------------14

**Supplementary Table 3A.** Banff g lesions by ptc lesions-------------------------------------	15 - 16

**Supplementary Table 3B.** Banff g lesions by i lesions----------------------------------------	17 - 18

**Supplementary Table 3C.** Banff g lesions by t lesions---------------------------------------- 19 - 20

**Supplementary Table 3D.** Banff ptc lesions by i lesions--------------------------------------	21 - 22

**Supplementary Table 3E.** Banff ptc lesions by t lesions--------------------------------------	23 - 24

**Supplementary Table 3F.** Banff i lesions by t lesions-----------------------------------------	25 - 26 **Supplementary Table 4.** Relationship of Banff Acute Lesion Score by Urinary Cell

Three-Gene Signature Score in 354 Biopsy-Matched Urine Specimens---------------------27 - 30

The Means Procedure-------------------------------------------------------------------------27 - 28

The GLM Procedure--------------------------------------------------------------------------29 - 30

**Supplementary Table 5.** Relationship of Composite Banff Scores by Urinary Cell

Three-Gene Signature Score in 354 Biopsy-Matched Urine Specimens---------------------31 - 34

The Means Procedure-------------------------------------------------------------------------31 - 32

The GLM Procedure--------------------------------------------------------------------------33 - 34

**SUPPLEMENTARY METHODS**

**Study Cohort (Figure 1)**

We evaluated 354 kidney allograft biopsy specimens and their corresponding urine samples, obtained prospectively from a total of 302 kidney transplant recipients. The cohort was recruited under institutional review-board approved research protocols # 940202002786 (“Use of PCR to Evaluate Renal Allograft Status”) and # 1207012730 (“Development of Gene Expression Monitoring (GEM) Biobank to Study Non-Invasive Biomarkers that Diagnose and Anticipate Post-Transplant Complications”). All participants signed written informed consent documents, underwent kidney allograft biopsy procedures at NewYork Presbyterian/Weill Cornell Medicine, and received their standardized post-transplant clinical follow-up care at our institution.

Biopsy procedures were undertaken either for clinical indications, such as increases in serum creatinine, proteinuria or other evidence of graft dysfunction, or as part of scheduled surveillance protocol. Of the 354 biopsy-urine pairs, 330 urine specimens were collected on the day of biopsy, 22 samples were obtained within 24 hours after the biopsy procedure, and 2 samples were collected within 3 days of biopsy; all urine collections followed a uniform, prespecified protocol ^1^. The biopsies were performed using the Monopty Disposable Core Biopsy Instrument, 18G (gauge size) × 16 cm (needle length) with 22 mm penetration depth (Product #121816, BD), and 2 biopsy cores were obtained during each biopsy procedure.

Clinical data were extracted data from a prospectively maintained, IRB- approved research database and verified against electronic medical record to ensure accuracy. The conduct of the study adhered to the ethical principles outlined in the Declaration of Helsinki.

**Banff Pathology Assessment of Kidney Allograft Biopsies**

Ultrasound-guided percutaneous needle core biopsies of kidney allografts were performed at NewYork-Presbyterian Hospital/Weill Cornell Medicine. All specimens were processed using histopathology protocols and interpreted by experienced renal pathologists according to the Banff classification of renal allograft pathology ^2^. Formalin-fixed paraffin-embedded sections were stained with hematoxylin and eosin, periodic acid-Schiff , and trichrome, and additional immunochemical studies for C4d and SV40 large T antigen were carried out on frozen or paraffin sections, as appropriate.

Banff acute lesion scores (0, 1, 2 or 3) were assigned semi-quantitatively for glomerulitis (g), peritubular capillaritis (ptc), interstitial inflammation (i), tubulitis (t), and intimal arteritis (v). Pathologists were masked to the urinary cell biomarker data at the time of biopsy interpretation. For analytic purposes, composite indices were derived by summing individual lesion scores to generate measures of microvascular inflammation (g + ptc) and tubulointerstitial inflammation (i+t), consistent with contemporary Banff guidance.

**Urinary Cell mRNA Quantification By Customized RT-qPCR**

Urine specimens, typically 50 mL of urine, were collected in sterile containers from kidney allograft recipients at the time of a clinically indicated or protocol surveillance biopsy and sent to the Gene Expression Monitoring (GEM) at Weill Cornell Medicine for urinary cell mRNA analysis.

Urine samples were centrifuged at 2000g for 30 min at room temperature to pellet the cellular fraction. The resulting urinary cell pellets were lysed in RNalater (50 μL), RLT buffer and beta-mercaptoethanol (350 μL), and RNA was extracted with the RNeasy mini kit (Qiagen, Cat. 74104). RNA concentration and purity were assessed using a NanoDrop spectrophotometer (ThermoFisher Scientific) using A260 and A260/A280 measurements.

Complementary DNA was synthesized from 1.0 μg in 100 μL (concentration) total RNA using the TaqMan reverse transcription kit (Applied Biosystems Cat. N808–0234). Reverse transcription was carried out in a 100 μL reaction containing 1× TaqMan RT buffer, 500 μM of each 4 dNTP, 2.5 μM of random hexamer, 0.4 Unit/μL of RNase inhibitor, 1.25 Unit/μL of MultiScribe Reverse Transcriptase and 5.5 mM MgCl2. The thermal profile consisted of 25°C for 10 min, 48°C for 30 min, and 95°C for 5 min.

We designed gene-specific oligonucleotide primers and hydrolysis TaqMan fluorogenic probes in our laboratory to quantify mRNA transcripts for CD3E, CXCL10, TGF-B1 and 18S rRNA. TGF-B1 mRNA used as a quality-control transcript for the customized RT-qPCR assays and 18S rRNA served both as a quality-control marker and as one of the components of the three-gene signature, as previously reported ^1^. Primer and probe sequences, as well as genomic coordinates, have been reported elsewhere ^1^.

Because urinary cells yield limited amount of total RNA, we applied a preamplification step before qPCR, as established in our laboratory ^1^. Each 10.0 μL preamplification reaction consisted of 3.0 μL cDNA, 5.0 μL Platinum® Multiplex PCR Master Mix, 1.68 μL primer mix containing 50 μM sense and 50 μM antisense primers and 0.32 μL water. Amplification was performed on a Veriti thermal cycler (Applied Biosystems) with 95°C for 2 min, followed by11 cycles of 95°C denaturing for 30s, 60°C for 90s for annealing, and 72°C for 1 min for primer extension, with a final extension at 72°C for 10 min. The amplified product was then diluted with 290 μL TE buffer, and 2.5 μL of the diluted sample was used for quantitative RT- qPCR.

Absolute transcript number was derived from a standard curve generated with an in-house synthesized Bak amplicon. The Bak standard was prepared at 1x10^7^copies/μL and serially diluted over six orders of magnitude the stock solution was diluted over 6 orders of magnitude to yield 1,000,000, 100,000, 10,000, 1000, 100, and 10 copies per μL. Each dilution was loaded in duplicate, amplified with Bak-specific primers and TaqMan probe, and used to generate a standard curve by plotting threshold cycle values for log of input copy number. This calibration defined the assay range, extending from 25 to 2.5 million copies.

The three-gene signature score was computed using the locked regression equation from the CTOT-04 study: Three-gene score = -6.1487+ 0.8534 log10 (CD3E /18S) + 0.6376 log10 (CXCL10/18S) + 1.6464 log10(18S) where the units of measurement in the customized RT-qPCR assays were copy numbers per microgram of total RNA isolated from the urinary cell pellets, and the units for 18S rRNA were number of copies (x10^-6^) per microgram of RNA of total RNA. In the equation, -6.1487 is the intercept, and 0.8534, 0.6376, and 1.6487 are the slopes (coefficients), respectively, for the log10 (CD3E /18S), log10 (CXCL10/18S) and log10(18S) values in the best -fitting logistic regression model. The intercept and slopes have no intrinsic units of measurement. This locked model allows score calculation to remain consistent across studies and avoids dataset-specific refitting.

**Rationale and strengths of statistical testing**

***Association among Banff acute lesion scores (Supplementary Tables 2 & 3A-F):*** The Banff acute lesion scores (g, ptc, i, t) are recorded on ordered categorical scales (ordinal scales) and are thus not treated as continuous variables. Accordingly, we used rank-based measures of association that operate on contingency tables rather than Pearson correlations or linear model, which assume interval-scale data and linearity. Goodman-Kruskal’s gamma and Somers’ D summarize the balance of concordant and discordant pairs across the cross-classified lesion scores, thereby quantifying whether higher grades of one lesion tend to accompany higher grades of another on standardized score from -1 to 1.

***Goodman–Kruskal’s gamma:*** It is a symmetric index of ordinal association and does not distinguish between predictor and outcome. This symmetry is advantageous when the goal is to determine whether two Banff lesions vary together, rather than to assign a directional causal interpretation. Gamma is defined by the difference between the proportions of concordant and discordant pairs among non-tied observations, which makes it relatively sensitive to ordered trends in sparse or skewed tables and provides an intuitive “proportional reduction in error” interpretation.

***Somers’ D:*** It is an asymmetric extension of these concordance-based measures and explicitly designates one variable as independent and the other as dependent. In situations where one Banff score is treated as the explanatory variable, and another as the response, Somers’ D therefore is more appropriate than gamma. By incorporating ties on the predictor, Somers’ D yields a more conservative estimate of associations when many biopsies share the same grade, and its definition parallels measures used to summarize predictive performance in ordinal regression, allowing coherence between exploratory correlation analyses and subsequent modeling.

***Use of contingency tables and cross‑tabulations****:* Estimating gamma and Somers’ D from contingency tables allowed us to keep the original ordinal categories, avoid imposing linearity or equal spacing of grades, and visually examine the joint distribution of lesion scores. The cross-tabulations presented in Supplementary Table 1 provide an easily interpretable basis for demonstrating that acute Banff lesions are positively and monotonically related across g, ptc, i, and t, while the associated gamma and Somers’ D place the strength and direction of these relationships on a common unitless scale.

***Rationale for Kruskal-Wallis testing (Figures 2 & 3) and Eta from one-way ANOVA (Table 1):*** For each Banff acute lesion category, we report the mean urinary cell three-gene score and corresponding confidence interval. These summaries give a clinically intuitive view of central tendency and precision for the continuous urinary cell three-gene biomarker across increasing lesion grades and make the magnitude of between -group differences directly interpretable on the original scale.

***Kruskal–Wallis test:*** To compare the urinary cell three-gene score across ordered Banff categories, we used Kruskal-Wallis test, a rank-based, distribution-free alternative to one-way ANOVA that does not require normal residuals or equal variances. This test is well-suited to our data, where the biomarker distribution is skewed and some lesion categories contain relatively few biopsies, and it addresses the global question of whether at least one category differs systematically in its biomarker distribution from others.

***Eta-squared (η²) from one‑way ANOVA****:* Eta (η) and Eta-squared (η²) from one‑way ANOVA were used as effect-size indices, quantifying the fractions of total variance in the three-gene score attributable to between-group differences across Banff acute lesion categories. In view of its close relationship to the coefficient of determination, η² provides a complementary, model-based summary of associations strength that is readily interpretable in terms of small, moderate , or large effects and enables comparison with other covariates and with external reports, even though our primary hypothesis testing was conducted with non-parametric Kruskal-Wallis procedure.

We depicted the distribution of the urinary cell three-gene signature values across Banff lesion scores with box-and-whisker plots. In these plots, the box boundaries represent the 25th and 75th percentiles, the horizontal line within each box represents the median, and the whiskers extend the conventional inner fences, defined as 1.5 times the interquartile range below the 25th percentile and above the 75th percentile.

***Strengths of combining these elements:*** Incorporating summary means and confidence intervals together with the Kruskal-Wallis test and η² permits concurrent evaluation of statistical significance, magnitude of effect, and potential clinical importance of the association between Banff acute lesion grade and the urinary cell three-gene signature. Taken together, these complementary metrices acknowledge the ordered, categorical nature of the Banff scores, remain robust in the presence of non-Gaussian biomarker distribution, and provide transparent, readily interpretable quantitative summaries that frame and corroborate the primary modeling findings presented in Supplemental Tables 3 and 4.

***Justification for general linear modeling of the urinary cell three-gene signature score (Supplementary Tables 4 & 5):*** We used a general linear modeling strategy to move beyond global, rank-based tests to obtain effect-size-oriented, model-based estimates of each histologic lesion’s independent association with the urinary cell three-gene score. In this framework, the three-gene signature (continuous outcome) is expressed as a linear function of Banff lesion categories and other covariates, with one-way ANOVA representing the special case in which only categorical predictors are included. Specifying each Banff lesion score as a categorical predictor allows estimation of adjusted mean difference in the biomarker across lesion levels while simultaneously accounting for the presence of other lesions- an analysis that cannot be performed using single-factor nonparametric procedures.

***Relative advantages over nonparametric tests:*** While Kruskal–Wallis and related rank-based procedures offer robust omnibus tests for distributional differences across Banff categories, they do not provide lesion-specific effect estimates. In contrast, general linear models yield regression coefficients or adjusted means that characterize both the direction and magnitude of the association between each individual lesion and the urinary cell three-gene score, accompanied by confidence intervals and P-values. This parametric, model-based strategy clarifies the independent contribution of each lesion, permits evaluation of collinearity and overlap among histologic variables, and creates a straightforward pathway to more elaborate multivariable and predictive modeling frameworks.

***Advantages of treating Banff scores as categorical predictors:*** Treating Banff lesion scores as categorical predictors rather than as a single numeric scale acknowledges that these grades represent ordered groups without assuming proportional spacing or a strictly linear relation with the biomarker. Within the general linear modeling framework, this parameterization allows flexible estimation of category-specific mean three-gene scores, accommodates nonmonotonic or irregular patterns across grades, and avoids imposing implausible linearity constraints. In addition, expressing effects as contrasts between clinically familiar categories (for examples, g0, g1, g2, g3) makes the resulting estimates directly interpretable to clinicians and pathologists who routinely use these lesion grades in practice.

***Complementarity with the nonparametric framework:*** Positioning the general linear models within this analysis provides a parametric counterpart to the rank-based nonparametric tests. The GLM framework yields adjusted, lesion -specific estimates of the association between each acute lesion score and the urinary cell three-gene signature, both when lesions are examined individually and when they are included concurrently in multivariable models. Taken together with the Kruskal-Wallis results, these model-based estimates reinforce the linkage between the three-gene signature score and histologic activity and demonstrate that the observed relationship are not artifacts of a single dominant lesion or of distributional features captured only by nonparametric methods.

***Rationale for proportional‑odds ordinal logistic models (Tables 2A & 4A-C):*** We applied proportional-odds ordinal logistic regression to evaluate whether the urinary cell three-gene signature predicts Banff acute lesion severity in a graded, statistically coherent manner that respects the ordered nature of the histologic scores. This modeling framework links the three-gene score to the cumulative odds of being in higher versus lower lesion categories, providing an unfed approach to quantify both overall and threshold -specific effects on ordered outcomes.

Because Banff lesion scores (0-3) constitute ordered categories rather than continuous measurements, the proportional-odds model is natural choice for characterizing how increments in the three-gene score shift the probability distribution across lesion grades. Fitting separate proportional-odds models for individual lesions and for composite Banff indices allows the biomarker’s progressive performance to be examined across microvascular and tubulointerstitial domains, while yielding a single summary odds ratio for each model when the proportional-odds assumption is reasonably satisfied.

***Strengths of the graded probability plots and odds ratios:*** Plots of model-based predicted probabilities against the urinary cell three-gene score recast the regression output into clinically intuitive “risk curves” making it apparent that increasing molecular signal corresponds to arising probability of more severe Banff lesions, with particularly pronounced gradients for the i and t components where the probability trajectories are steepest. The accompanying odds ratios and likelihood-ratio chi-squares statistics summarize, respectively, the per-unit change in the odds of more advanced histologic injury and the overall incremental information contributed by the biomarker, thereby demonstrating that the urinary three-gene signature explains a substantial proportion of variability in lesion severity across both individual lesions and composite Banff scores.

***Rationale and strengths of calibration and assumption checks:*** Evaluating calibration with the Hosmer-Lemeshow goodness-of-fit allows comparison of predicted probabilities from the ordinal logistic models with the observed frequencies of each Banff lesion category, thereby supporting the reliability of the risk estimates for most endpoints and explicitly highlighting modest miscalibration for the combined i+t score was present. Systematic testing of the proportional-odds assumption further differentiates lesions for which a single summary odds ration provides an adequate description of the association between the three-gene signature and lesion severity (g, ptc, i, g+ptc) from those in which the biomarker-severity relationship varies across cut points, helping to avoid over-simplified inference and to guide flexible modeling or threshold-specific estimates are warranted.

***Rationale for generalized (multinomial) logistic models (Table 2B & 4D):*** When the proportional-odds assumption was not tenable for the t lesion and composite Banff scores that incorporate t, we replaced the ordinal proportionate-odds model with a generalized (multinomial) logistic specification to obtain category specific effects of the three-gene signature on lesion severity.

This more flexible nominal model demonstrates a marked decline in the probability of t=0 and a corresponding increase in the probability of t=3 as the three-gene score rises, underscoring that tabulated and graphic summaries respect the intrinsically non-proportional patter in the data.

***Overall strengths of this modeling strategy:*** Taken together, the proportional-odds logistic models and the generalized (multinomial) logistic models provide a unified framework for appraising the urinary cell three-gene signature as a noninvasive diagnostic biomarker. This strategy preserves the inherent ordering of Banff acute lesion grades, captures strong monotonic relationships where proportionality is reasonable, explicitly documents model fit and key assumptions, and switches to more flexible multinomial model specifications when the proportional-odds constraint is untenable. By combining rigorous model checking with probability and odds-ratio summaries that map directly onto clinically meaningful rejection lesions, the analysis yields risk gradients that are both statistically defensible and readily interpretable at the bedside.

**SUPPLEMENTARY TABLE 1A. Demographics and Clinical Characteristics of Adult Kidney Allograft Recipients.**

| **Characteristics** | **Kidney Transplant Study groups** | | |
| --- | --- | --- | --- |
|  | Acute Rejection (AR) | No Rejection (NR) | P value^a^ |
| Kidney allograft recipients, N^b^ | 166 | 136 |  |
| Kidney allograft biopsies, N^c^ | 205 | 149 |  |
| Biopsy matched urine samples, N | 205 | 149 |  |
| **Kidney recipient information** |  |  |  |
| Age, yr, mean (SD) | 49 (16) | 49 (16) | 0.7231 |
| Women, N (%) | 79 (47) | 49 (36) | 0.0472 |
| Racial categories |  |  |  |
| White, N (%) | 59 (29) | 49 (36) | 0.0689 |
| Black, N (%) | 46 (22) | 38 (28) |  |
| Other, N (%) | 100 (49) | 49 (36) |  |
| **Cause of native kidney disease** |  |  |  |
| Diabetes mellitus, N (%) | 31 (19) | 39 (29) |  |
| Glomerular disease, N (%) | 41 (25) | 34 (25) |  |
| Hypertension, N (%) | 47 (28) | 33 (24) | 0.2581 |
| Lupus nephritis, N (%) | 8 (5) | 2 (1) |  |
| Polycystic disease, N (%) | 13 (8) | 8 (6) |  |
| Other, N (%) | 26 (16) | 20 (15) |  |
| **Type of Kidney Allograft Transplant** |  |  | 0.9664 |
| Living Related, N (%) | 42 (25) | 33 (24) |  |
| Living Unrelated, N (%) | 56 (34) | 45 (33) |  |
| Deceased, N (%) | 68 (41) | 58 (43) |  |
| **Induction therapy** |  |  | 0.8927 |
| Antithymocyte globulin, N (%) | 133 (80) | 106 (78) |  |
| Basiliximab, N (%) | 19 (11) | 17 (13) |  |
| Other, N(%) | 14 (8) | 13 (10) |  |
| **Maintenance therapy** |  |  | 0.279 |
| Tacrolimus, N (%) | 155 (93) | 127 (93) |  |
| Mycophenolate mofetil, N (%) | 153 (92) | 127 (93) |  |
| Corticosteroid maintenance, N (%) | 42 (25) | 20 (15) |  |
| Other, N (%) | 11 (7) | 9 (7) |  |
| **Reason for Biopsy** |  |  |  |
| Clinically indicated, N (%) | 170 (83) | 118 (79) | 0.4081 |
| Surveillance, N (%) | 35 (17) | 31 (21) |  |
| Time from transplantation to biopsy, Days, median (IQR) | 373 (104-1211) | 181 (70-1362) | 0.1824 |
| Clinically indicated biopsies, Days, median (IQR) | 359 (94-1302) | 151 (58-1611) | 0.2739 |
| Surveillance biopsies, Days, median (IQR) | 399 (113-547) | 268 (111-478) | 0.4883 |
| Serum creatinine at index biopsy, mg/dl, median (IQR) | 2.01 (1.40-3.06) | 1.85 (1.45-3.06) | 0.9142 |
| **Donor Specific Antibodies (DSA) at index allograft biopsy**^d^ |  |  |  |
| Data available, N (%) | 162 (79) | 121 (81) | 0.6872 |
| DSA present, N (%) | 65 (40) | 19 (16) | 0.0003 |
| DSA against HLA Class I, N (%) | 11 (5) | 5 (4) |  |
| DSA against HLA Class II, N (%) | 33 (20) | 13 (9) |  |
| DSA against HLA Class I & II, N (%) | | 21 (10) | 1 (1) |

Continuous variables are summarized as mean (standard deviation) or median (interquartile range), and categorical variables as number (percentage).

a: Two-sided P values were calculated using Mann-Whitney U test for continuous variables and Fisher's exact test for categorical values.

b: Among the 166 kidney allograft recipients in the AR group, 133 had one biopsy, and 33 had two or more biopsies. Among the 136 kidney allograft recipients in the NR group, 125 had one biopsy and 11 had two or more biopsies. In the AR biopsy subtypes included: Antibody mediated rejection (n= 61), T-cell mediated rejection (n=41), Borderline rejection (n=27), Mixed rejection (n=23), and Microvascular rejection (n=27). Among the 166 kidney allograft recipients in the AR group, 133 had one biopsy, and 33 had two or more biopsies. Among the 136 kidney allograft recipients in the NR group, 125 had one biopsy and 11 had two or more biopsies.

c: Fourteen of the 136 kidney allograft recipients in the NR group had a previous AR biopsy in the AR group.The interval between the AR and subsequent NR biopsy was 10 months in 1 recipient, 9 months in 2 recipients, 6 months in 1 recipient, 5 months in 1 recipient, 4 months in 6 recipients, 2 months in 1 recipient, and 1 month in 2 recipients.

d: DSA were detected in kidney allograft recipients using LABScreen Single Antigen (LSA) assays - LS1A04 for class I and LS2A01 for class II antibodies - performed in a clinical histocompatibility laboratory. A DSA was considered positive if the mean fluorescence intensity (MFI) exceeded 2000.

**SUPPLEMENTARY TABLE 1B. Allograft Biopsy Histopathology Characteristics.**

| **Characteristics** | **Kidney Transplant Study groups** | | | | |
| --- | --- | --- | --- | --- | --- |
|  | Acute Rejection (AR) | | No Rejection (NR) | | P value^a^ |
| **Kidney allograft biopsy findings**^b^ | **median (IQR)** | **mean (SD)** | **median (IQR)** | **mean (SD)** |  |
| Number of glomeruli | 16 (12-25) | 19 (11) | 18 (12-25) | 19.8 (10.7) | 0.4276 |
| Banff acute lesion scores |  |  |  |  |  |
| Tubulitis (t) score | 1 (0-2) | 1.10 (1.16) | 0 (0-0) | 0.10 (0.08) | <0.0001 |
| Interstitial inflammation (i) score | 1 (0-2) | 1.34 (1.15) | 0 (0-0) | 0.18 (0.40) | <0.0001 |
| Glomerulitis (g) score | 1 (0-1) | 0.92 (1.04) | 0 (0-0) | 0.04 (0.20) | <0.0001 |
| Peritubular capillaritis (ptc) score | 2 (1-3) | 1.82 (1.06) | 0 (0-0) | 0.11 (0.31) | <0.0001 |
| Vascular inflammation (v) score | 0 (0-0) | 0.11 (0.39) | 0 (0-0) | 0 (0) | 0.0005 |
| Banff chronic lesion scores |  |  |  |  |  |
| Tubular atrophy (ct) score | 1 (0-2) | 1.02 (1.02) | 0 (0-1) | 0.80 (1.01) | 0.0245 |
| Interstitial fibrosis (ci) score | 1 (0-2) | 0.97 (1.01) | 0 (0-1) | 0.65 (0.89) | 0.0023 |
| Chronic glomerulopathy (cg) score | 0 (0-0) | 0.50 (1.04) | 0 (0-0) | 0.04 (0.31) | <0.0001 |
| Vascular fibrosis intimal thickening (cv) score | 2 (1-2) | 1.60 (1.01) | 0 (1-2) | 1.35 (1.07) | 0.031 |
| Arteriolar hyalinosis (ah) score | 0 (0-2) | 0.78 (1.15) | 0 (0-1) | 0.79 (1.09) | 0.7388 |
| Positive staining for SV40 large T antigen, N (%) | 0 (0) | | 0 (0) | | >0.9999 |
| Positive staining for complement component 4d, N (%) | 20 (10) | | 3 (2) | | 0.0013 |

a: P values were calculated using Mann-Whitney U test for continuous variables and Fisher's exact test for categorical values.

b: Kidney allograft biopsies were classified according to the Banff 2022 update of the Banff '97 classification system.

**SUPPLEMENTARY TABLE 1C. Urinary Cell RNA Characteristics.**

| **Characteristics** | **Kidney Transplant Study groups** | | |
| --- | --- | --- | --- |
|  | Acute Rejection (AR) | No Rejection (NR) | P value^a^ |
| **Urinary cell RNA**^b^ |  |  |  |
| Total RNA, ug, median (IQR) | 0.97 (0.48-2.16) | 0.98 (0.48-1.85) | 0.5011 |
| Total RNA purity, A260/A280, median (IQR) | 1.94 (1.90-2.00) | 1.94 (1.88-2.01) | 0.3684 |

a: P values were calculated using Mann-Whitney U test.

b: Total RNA from biopsy matched urine samples were isolated using RNeasy Mini Kit. The total RNA yield (absorbance at 260) and quality (A260/280 ratio) were measured using Nanodrop spectrophotometer.

**SUPPLEMENTARY TABLE 2.** **Summary Statistics Crosstabulations for each pair of lesion scores.**

|  |  | ptc lesion | i lesion | t lesion |
| --- | --- | --- | --- | --- |
| g lesion score | chi-square | 146.6 | 41.9 | 19.8 |
|  | gamma | 0.709 | 0.340 | 0.325 |
|  | Somers’ D Col\|Row | 0.567 | 0.235 | 0.186 |
|  | Somers’ D Row\|Col | 0.416 | 0.187 | 0.185 |
| ptc lesion score | chi-square |  | 89.6 | 80.5 |
|  | gamma |  | 0.556 | 0.590 |
|  | Somers’ D Col\|Row |  | 0.386 | 0.341 |
|  | Somers’ D Row\|Col |  | 0.418 | 0.462 |
| i lesion score | chi-square |  |  | 424.1 |
|  | gamma |  |  | 0.969 |
|  | Somers’ D Col\|Row |  |  | 0.702 |
|  | Somers’ D Row\|Col |  |  | 0.874 |

Gamma: a measure correlation between two ordinal variables (treats rows and columns symmetrically)

Somers’ D, Col|Row: The probability that the Column variable increases when the Row variable increases

Somers’ D, Row|Col: The probability that the Row variable increases when the Column variable increases

**SUPPLEMENTARY TABLE 3A. Banff g lesions by ptc lesions.**

***Crosstabulations of Banff acute lesion scores in 354 kidney allograft biopsies included in this study***

***The FREQ Procedure***

```
                              g_lesions(g - Glomerulitis(0, 1, 2, 3))
                              ptc_lesions(Peritubular capillaritis(0, 1, 2, 3))

                               Frequency|
                               Percent  |
                               Row Pct  |
                               Col Pct  |       0|       1|       2|       3|  Total
                               ---------+--------+--------+--------+--------+
                                      0 |    143 |     50 |     22 |     21 |    236
                                        |  40.40 |  14.12 |   6.21 |   5.93 |  66.67
                                        |  60.59 |  21.19 |   9.32 |   8.90 |
                                        |  88.82 |  72.46 |  42.31 |  29.17 |
                               ---------+--------+--------+--------+--------+
                                      1 |     13 |     17 |     23 |     15 |     68
                                        |   3.67 |   4.80 |   6.50 |   4.24 |  19.21
                                        |  19.12 |  25.00 |  33.82 |  22.06 |
                                        |   8.07 |  24.64 |  44.23 |  20.83 |
                               ---------+--------+--------+--------+--------+
                                      2 |      2 |      1 |      4 |     16 |     23
                                        |   0.56 |   0.28 |   1.13 |   4.52 |   6.50
                                        |   8.70 |   4.35 |  17.39 |  69.57 |
                                        |   1.24 |   1.45 |   7.69 |  22.22 |
                               ---------+--------+--------+--------+--------+
                                      3 |      3 |      1 |      3 |     20 |     27
                                        |   0.85 |   0.28 |   0.85 |   5.65 |   7.63
                                        |  11.11 |   3.70 |  11.11 |  74.07 |
                                        |   1.86 |   1.45 |   5.77 |  27.78 |
                               ---------+--------+--------+--------+--------+
                               Total         161       69       52       72      354
                                           45.48    19.49    14.69    20.34   100.00

                                  Statistics for Table of g lesions by ptc_lesions

                               Statistic                     DF       Value      Prob
                               ------------------------------------------------------
                               Chi-Square                     9    146.6102    <.0001
                               Likelihood Ratio Chi-Square    9    131.7267    <.0001
                               Mantel-Haenszel Chi-Square     1    104.5670    <.0001
                               Phi Coefficient                       0.6435
                               Contingency Coefficient               0.5412
                               Cramer's V                            0.3716

                                                                                     95%
                    Statistic                              Value       ASE     Confidence Limits
                    ----------------------------------------------------------------------------
                    Gamma                                 0.7089    0.0459     0.6190     0.7988
                    Kendall's Tau-b                       0.4857    0.0404     0.4066     0.5648
                    Stuart's Tau-c                        0.3843    0.0363     0.3130     0.4555

                    Somers' D C|R                         0.5667    0.0453     0.4779     0.6554
                    Somers' D R|C                         0.4164    0.0385     0.3409     0.4918

                    Pearson Correlation                   0.5443    0.0447     0.4566     0.6319
                    Spearman Correlation                  0.5331    0.0441     0.4467     0.6194

                    Lambda Asymmetric C|R                 0.2124    0.0403     0.1334     0.2915
                    Lambda Asymmetric R|C                 0.0085    0.0566     0.0000     0.1194
                    Lambda Symmetric                      0.1350    0.0386     0.0595     0.2106

                    Uncertainty Coefficient C|R           0.1450    0.0241     0.0978     0.1923
                    Uncertainty Coefficient R|C           0.1936    0.0304     0.1340     0.2532
                    Uncertainty Coefficient Symmetric     0.1658    0.0267     0.1135     0.2182

                                                 Sample Size = 354
```

**SUPPLEMENTARY TABLE 3B. Banff g lesions by i lesions.**

```
                               g_lesions(g - Glomerulitis (0, 1, 2, 3))
                               i_lesions(i - Interstitial Inflammation (0, 1, 2,3))

                               Frequency|
                               Percent  |
                               Row Pct  |
                               Col Pct  |       0|       1|       2|       3|  Total
                               ---------+--------+--------+--------+--------+
                                      0 |    141 |     54 |     18 |     23 |    236
                                        |  39.83 |  15.25 |   5.08 |   6.50 |  66.67
                                        |  59.75 |  22.88 |   7.63 |   9.75 |
                                        |  75.81 |  63.53 |  54.55 |  46.00 |
                               ---------+--------+--------+--------+--------+
                                      1 |     33 |      8 |      9 |     18 |     68
                                        |   9.32 |   2.26 |   2.54 |   5.08 |  19.21
                                        |  48.53 |  11.76 |  13.24 |  26.47 |
                                        |  17.74 |   9.41 |  27.27 |  36.00 |
                               ---------+--------+--------+--------+--------+
                                      2 |      7 |      8 |      4 |      4 |     23
                                        |   1.98 |   2.26 |   1.13 |   1.13 |   6.50
                                        |  30.43 |  34.78 |  17.39 |  17.39 |
                                        |   3.76 |   9.41 |  12.12 |   8.00 |
                               ---------+--------+--------+--------+--------+
                                      3 |      5 |     15 |      2 |      5 |     27
                                        |   1.41 |   4.24 |   0.56 |   1.41 |   7.63
                                        |  18.52 |  55.56 |   7.41 |  18.52 |
                                        |   2.69 |  17.65 |   6.06 |  10.00 |
                               ---------+--------+--------+--------+--------+
                               Total         186       85       33       50      354
                                           52.54    24.01     9.32    14.12   100.00

                                   Statistics for Table of g_lesions by i_lesions

                               Statistic                     DF       Value      Prob
                               ------------------------------------------------------
                               Chi-Square                     9     41.9423    <.0001
                               Likelihood Ratio Chi-Square    9     40.4604    <.0001
                               Mantel-Haenszel Chi-Square     1     15.1424    <.0001
                               Phi Coefficient                       0.3442
                               Contingency Coefficient               0.3255
                               Cramer's V                            0.1987

                                                                                     95%
                    Statistic                              Value       ASE     Confidence Limits
                    ----------------------------------------------------------------------------
                    Gamma                                 0.3405    0.0651     0.2129     0.4681
                    Kendall's Tau-b                       0.2097    0.0439     0.1236     0.2959
                    Stuart's Tau-c                        0.1593    0.0341     0.0925     0.2260

                    Somers' D C|R                         0.2348    0.0491     0.1387     0.3310
                    Somers' D R|C                         0.1873    0.0400     0.1089     0.2657

                    Pearson Correlation                   0.2071    0.0488     0.1114     0.3028
                    Spearman Correlation                  0.2401    0.0506     0.1410     0.3392

                    Lambda Asymmetric C|R                 0.0655    0.0340     0.0000     0.1322
                    Lambda Asymmetric R|C                 0.0000    0.0000     0.0000     0.0000
                    Lambda Symmetric                      0.0385    0.0201     0.0000     0.0778

                    Uncertainty Coefficient C|R           0.0485    0.0150     0.0192     0.0778
                    Uncertainty Coefficient R|C           0.0595    0.0180     0.0242     0.0948
                    Uncertainty Coefficient Symmetric     0.0534    0.0163     0.0215     0.0854

                                                 Sample Size = 354
```

**SUPPLEMENTARY TABLE 3C. Banf g lesions by t lesion**

```
                               g_lesions(g - Glomerulitis (0, 1, 2, 3))
                               t_lesions(t - Tubulitis  (0,1,2,3))

                               Frequency|
                               Percent  |
                               Row Pct  |
                               Col Pct  |       0|       1|       2|       3|  Total
                               ---------+--------+--------+--------+--------+
                                      0 |    175 |     24 |     17 |     20 |    236
                                        |  49.44 |   6.78 |   4.80 |   5.65 |  66.67
                                        |  74.15 |  10.17 |   7.20 |   8.47 |
                                        |  73.53 |  54.55 |  51.52 |  51.28 |
                               ---------+--------+--------+--------+--------+
                                      1 |     38 |      9 |     10 |     11 |     68
                                        |  10.73 |   2.54 |   2.82 |   3.11 |  19.21
                                        |  55.88 |  13.24 |  14.71 |  16.18 |
                                        |  15.97 |  20.45 |  30.30 |  28.21 |
                               ---------+--------+--------+--------+--------+
                                      2 |     11 |      4 |      4 |      4 |     23
                                        |   3.11 |   1.13 |   1.13 |   1.13 |   6.50
                                        |  47.83 |  17.39 |  17.39 |  17.39 |
                                        |   4.62 |   9.09 |  12.12 |  10.26 |
                               ---------+--------+--------+--------+--------+
                                      3 |     14 |      7 |      2 |      4 |     27
                                        |   3.95 |   1.98 |   0.56 |   1.13 |   7.63
                                        |  51.85 |  25.93 |   7.41 |  14.81 |
                                        |   5.88 |  15.91 |   6.06 |  10.26 |
                               ---------+--------+--------+--------+--------+
                               Total         238       44       33       39      354
                                           67.23    12.43     9.32    11.02   100.00

                                   Statistics for Table of g_lesions by t_lesions

                               Statistic                     DF       Value      Prob
                               ------------------------------------------------------
                               Chi-Square                     9     19.8331    0.0190
                               Likelihood Ratio Chi-Square    9     18.3521    0.0313
                               Mantel-Haenszel Chi-Square     1      9.1154    0.0025
                               Phi Coefficient                       0.2367
                               Contingency Coefficient               0.2303
                               Cramer's V                            0.1367

                                                                                     95%
                    Statistic                              Value       ASE     Confidence Limits
                    ----------------------------------------------------------------------------
                    Gamma                                 0.3247    0.0753     0.1772     0.4722
                    Kendall's Tau-b                       0.1860    0.0486     0.0908     0.2812
                    Stuart's Tau-c                        0.1265    0.0336     0.0606     0.1924

                    Somers' D C|R                         0.1865    0.0491     0.0903     0.2827
                    Somers' D R|C                         0.1854    0.0488     0.0897     0.2811

                    Pearson Correlation                   0.1607    0.0542     0.0544     0.2670
                    Spearman Correlation                  0.2063    0.0539     0.1007     0.3120

                    Lambda Asymmetric C|R                 0.0000    0.0000     0.0000     0.0000
                    Lambda Asymmetric R|C                 0.0000    0.0000     0.0000     0.0000
                    Lambda Symmetric                      0.0000    0.0000     0.0000     0.0000

                    Uncertainty Coefficient C|R           0.0262    0.0125     0.0017     0.0506
                    Uncertainty Coefficient R|C           0.0270    0.0128     0.0018     0.0521
                    Uncertainty Coefficient Symmetric     0.0266    0.0126     0.0018     0.0513

                                                 Sample Size = 354
```

**SUPPLEMENTARY TABLE 3D. Banff ptc lesions by i lesion**

```
                               ptc_lesions(ptc inflammation (0, 1, 2, 3))
                               i_lesions(i - Interstitial Inflammation (0, 1, 2,3))

                               Frequency|
                               Percent  |
                               Row Pct  |
                               Col Pct  |       0|       1|       2|       3|  Total
                               ---------+--------+--------+--------+--------+
                                      0 |    116 |     38 |      5 |      2 |    161
                                        |  32.77 |  10.73 |   1.41 |   0.56 |  45.48
                                        |  72.05 |  23.60 |   3.11 |   1.24 |
                                        |  62.37 |  44.71 |  15.15 |   4.00 |
                               ---------+--------+--------+--------+--------+
                                      1 |     33 |     15 |      9 |     12 |     69
                                        |   9.32 |   4.24 |   2.54 |   3.39 |  19.49
                                        |  47.83 |  21.74 |  13.04 |  17.39 |
                                        |  17.74 |  17.65 |  27.27 |  24.00 |
                               ---------+--------+--------+--------+--------+
                                      2 |     24 |      9 |      9 |     10 |     52
                                        |   6.78 |   2.54 |   2.54 |   2.82 |  14.69
                                        |  46.15 |  17.31 |  17.31 |  19.23 |
                                        |  12.90 |  10.59 |  27.27 |  20.00 |
                               ---------+--------+--------+--------+--------+
                                      3 |     13 |     23 |     10 |     26 |     72
                                        |   3.67 |   6.50 |   2.82 |   7.34 |  20.34
                                        |  18.06 |  31.94 |  13.89 |  36.11 |
                                        |   6.99 |  27.06 |  30.30 |  52.00 |
                               ---------+--------+--------+--------+--------+
                               Total         186       85       33       50      354
                                           52.54    24.01     9.32    14.12   100.00

Statistics for Table of ptc_lesions by i_lesions

                               Statistic                     DF       Value      Prob
                               ------------------------------------------------------
                               Chi-Square                     9     89.5818    <.0001
                               Likelihood Ratio Chi-Square    9    100.4215    <.0001
                               Mantel-Haenszel Chi-Square     1     79.9619    <.0001
                               Phi Coefficient                       0.5030
                               Contingency Coefficient               0.4494
                               Cramer's V                            0.2904

                                                                                     95%

Statistic                              Value       ASE     Confidence Limits
                    ----------------------------------------------------------------------------
                    Gamma                                 0.5563    0.0458     0.4665     0.6461
                    Kendall's Tau-b                       0.4017    0.0376     0.3280     0.4754
                    Stuart's Tau-c                        0.3558    0.0344     0.2883     0.4233

                    Somers' D C|R                         0.3855    0.0376     0.3118     0.4593
                    Somers' D R|C                         0.4185    0.0384     0.3433     0.4938

                    Pearson Correlation                   0.4759    0.0421     0.3935     0.5584
                    Spearman Correlation                  0.4664    0.0429     0.3823     0.5506

                    Lambda Asymmetric C|R                 0.0774    0.0357     0.0074     0.1474
                    Lambda Asymmetric R|C                 0.1503    0.0313     0.0889     0.2116
                    Lambda Symmetric                      0.1163    0.0301     0.0574     0.1753

                    Uncertainty Coefficient C|R           0.1204    0.0198     0.0815     0.1593
                    Uncertainty Coefficient R|C           0.1106    0.0188     0.0737     0.1475
                    Uncertainty Coefficient Symmetric     0.1153    0.0193     0.0775     0.1530

                                                 Sample Size = 354
```

**SUPPLEMENTARY TABLE 3E. Banff ptc lesions by t lesion**

```
                               ptc_lesions(ptc inflammation (0, 1, 2, 3))
                               t_lesions(t - Tubulitis  (0,1,2,3))

                               Frequency|
                               Percent  |
                               Row Pct  |
                               Col Pct  |       0|       1|       2|       3|  Total
                               ---------+--------+--------+--------+--------+
                                      0 |    141 |     16 |      2 |      2 |    161
                                        |  39.83 |   4.52 |   0.56 |   0.56 |  45.48
                                        |  87.58 |   9.94 |   1.24 |   1.24 |
                                        |  59.24 |  36.36 |   6.06 |   5.13 |
                               ---------+--------+--------+--------+--------+
                                      1 |     42 |      7 |      9 |     11 |     69
                                        |  11.86 |   1.98 |   2.54 |   3.11 |  19.49
                                        |  60.87 |  10.14 |  13.04 |  15.94 |
                                        |  17.65 |  15.91 |  27.27 |  28.21 |
                               ---------+--------+--------+--------+--------+
                                      2 |     30 |      6 |      9 |      7 |     52
                                        |   8.47 |   1.69 |   2.54 |   1.98 |  14.69
                                        |  57.69 |  11.54 |  17.31 |  13.46 |
                                        |  12.61 |  13.64 |  27.27 |  17.95 |
                               ---------+--------+--------+--------+--------+
                                      3 |     25 |     15 |     13 |     19 |     72
                                        |   7.06 |   4.24 |   3.67 |   5.37 |  20.34
                                        |  34.72 |  20.83 |  18.06 |  26.39 |
                                        |  10.50 |  34.09 |  39.39 |  48.72 |
                               ---------+--------+--------+--------+--------+
                               Total         238       44       33       39      354
                                           67.23    12.43     9.32    11.02   100.00

                                  Statistics for Table of ptc_lesions by t_lesions

                               Statistic                     DF       Value      Prob
                               ------------------------------------------------------
                               Chi-Square                     9     80.5242    <.0001
                               Likelihood Ratio Chi-Square    9     90.2946    <.0001
                               Mantel-Haenszel Chi-Square     1     68.9667    <.0001
                               Phi Coefficient                       0.4769
                               Contingency Coefficient               0.4305
                               Cramer's V                            0.2754

                                                                                     95%
                    Statistic                              Value       ASE     Confidence Limits
                    ----------------------------------------------------------------------------
                    Gamma                                 0.5900    0.0482     0.4955     0.6844
                    Kendall's Tau-b                       0.3968    0.0384     0.3215     0.4721
                    Stuart's Tau-c                        0.3149    0.0329     0.2503     0.3794

                    Somers' D C|R                         0.3412    0.0361     0.2705     0.4119
                    Somers' D R|C                         0.4615    0.0434     0.3765     0.5465

                    Pearson Correlation                   0.4420    0.0440     0.3558     0.5282
                    Spearman Correlation                  0.4545    0.0435     0.3691     0.5398

                    Lambda Asymmetric C|R                 0.0000    0.0000     0.0000     0.0000
                    Lambda Asymmetric R|C                 0.1451    0.0287     0.0887     0.2014
                    Lambda Symmetric                      0.0906    0.0175     0.0563     0.1249

                    Uncertainty Coefficient C|R           0.1288    0.0222     0.0853     0.1723
                    Uncertainty Coefficient R|C           0.0994    0.0181     0.0640     0.1348
                    Uncertainty Coefficient Symmetric     0.1122    0.0198     0.0734     0.1510

                                                 Sample Size = 354
```

**SUPPLEMENTARY TABLE 3F. Banff i lesions by t lesion**

```
                               Supplemental Table 1F.i_lesions by t_lesions

                               i_lesions(i - Interstitial Inflammation (0, 1, 2,3))
                               t_lesions(t - Tubulitis  (0,1,2,3))

                               Frequency|
                               Percent  |
                               Row Pct  |
                               Col Pct  |       0|       1|       2|       3|  Total
                               ---------+--------+--------+--------+--------+
                                      0 |    184 |      2 |      0 |      0 |    186
                                        |  51.98 |   0.56 |   0.00 |   0.00 |  52.54
                                        |  98.92 |   1.08 |   0.00 |   0.00 |
                                        |  77.31 |   4.55 |   0.00 |   0.00 |
                               ---------+--------+--------+--------+--------+
                                      1 |     50 |     33 |      2 |      0 |     85
                                        |  14.12 |   9.32 |   0.56 |   0.00 |  24.01
                                        |  58.82 |  38.82 |   2.35 |   0.00 |
                                        |  21.01 |  75.00 |   6.06 |   0.00 |
                               ---------+--------+--------+--------+--------+
                                      2 |      3 |      6 |     18 |      6 |     33
                                        |   0.85 |   1.69 |   5.08 |   1.69 |   9.32
                                        |   9.09 |  18.18 |  54.55 |  18.18 |
                                        |   1.26 |  13.64 |  54.55 |  15.38 |
                               ---------+--------+--------+--------+--------+
                                      3 |      1 |      3 |     13 |     33 |     50
                                        |   0.28 |   0.85 |   3.67 |   9.32 |  14.12
                                        |   2.00 |   6.00 |  26.00 |  66.00 |
                                        |   0.42 |   6.82 |  39.39 |  84.62 |
                               ---------+--------+--------+--------+--------+
                               Total         238       44       33       39      354
                                           67.23    12.43     9.32    11.02   100.00

                                   Statistics for Table of i_lesions by t_lesions

                               Statistic                     DF       Value      Prob
                               ------------------------------------------------------
                               Chi-Square                     9    424.0775    <.0001
                               Likelihood Ratio Chi-Square    9    384.2356    <.0001
                               Mantel-Haenszel Chi-Square     1    272.6237    <.0001
                               Phi Coefficient                       1.0945
                               Contingency Coefficient               0.7383
                               Cramer's V                            0.6319

                                                                                     95%
                    Statistic                              Value       ASE     Confidence Limits
                    ----------------------------------------------------------------------------
                    Gamma                                 0.9692    0.0091     0.9512     0.9871
                    Kendall's Tau-b                       0.7831    0.0213     0.7415     0.8248
                    Stuart's Tau-c                        0.5964    0.0332     0.5313     0.6616

                    Somers' D C|R                         0.7015    0.0295     0.6438     0.7593
                    Somers' D R|C                         0.8742    0.0171     0.8407     0.9077

                    Pearson Correlation                   0.8788    0.0166     0.8464     0.9113
                    Spearman Correlation                  0.8271    0.0218     0.7843     0.8699

                    Lambda Asymmetric C|R                 0.4052    0.0493     0.3085     0.5018
                    Lambda Asymmetric R|C                 0.4881    0.0395     0.4107     0.5655
                    Lambda Symmetric                      0.4542    0.0367     0.3822     0.5262

                    Uncertainty Coefficient C|R           0.5480    0.0304     0.4885     0.6075
                    Uncertainty Coefficient R|C           0.4606    0.0303     0.4012     0.5199
                    Uncertainty Coefficient Symmetric     0.5005    0.0298     0.4421     0.5589

                                                 Sample Size = 354
```

**SUPPLEMENTARY TABLE 4. Relationship of Banff Acute Lesion Score by Urinary Cell Three-Gene Signature Score in**

**354 Biopsy-Matched Urine Specimens**

***The MEANS Procedure***

| **Analysis Variable : CTOT04_sig** | | | | | | | | | | |
| --- | --- | --- | --- | --- | --- | --- | --- | --- | --- | --- |
| **g Banff lesion score** | **N Obs** | **N** | **Minimum** | **Lower Quartile** | **Median** | **Upper Quartile** | **Maximum** | **Mean** | **Std Dev** | **Std Error** |
| 0 | 236 | 236 | -4.201 | -2.174 | -1.329 | -0.538 | 2.189 | -1.274 | 1.246 | 0.081 |
| 1 | 68 | 68 | -3.568 | -1.759 | -0.907 | 0.068 | 1.806 | -0.855 | 1.218 | 0.148 |
| 2 | 23 | 23 | -2.610 | -1.306 | -0.679 | 0.072 | 1.289 | -0.567 | 0.985 | 0.205 |
| 3 | 27 | 27 | -2.937 | -0.951 | 0.167 | 0.718 | 2.169 | -0.161 | 1.269 | 0.244 |

| **Analysis Variable : CTOT04_sig** | | | | | | | | | | |
| --- | --- | --- | --- | --- | --- | --- | --- | --- | --- | --- |
| **ptc Banff lesion score** | **N Obs** | **N** | **Minimum** | **Lower Quartile** | **Median** | **Upper Quartile** | **Maximum** | **Mean** | **Std Dev** | **Std Error** |
| 0 | 161 | 161 | -4.201 | -2.220 | -1.344 | -0.592 | 1.494 | -1.367 | 1.176 | 0.093 |
| 1 | 69 | 69 | -4.118 | -2.230 | -1.132 | -0.249 | 2.189 | -1.041 | 1.393 | 0.168 |
| 2 | 52 | 52 | -3.086 | -1.896 | -1.171 | -0.134 | 2.169 | -0.957 | 1.289 | 0.179 |
| 3 | 72 | 72 | -3.568 | -1.266 | -0.551 | 0.267 | 2.107 | -0.481 | 1.130 | 0.133 |

| **Analysis Variable : CTOT04_sig** | | | | | | | | | | |
| --- | --- | --- | --- | --- | --- | --- | --- | --- | --- | --- |
| **i Banff lesion score** | **N Obs** | **N** | **Minimum** | **Lower Quartile** | **Median** | **Upper Quartile** | **Maximum** | **Mean** | **Std Dev** | **Std Error** |
| 0 | 186 | 186 | -4.201 | -2.226 | -1.353 | -0.592 | 1.505 | -1.378 | 1.154 | 0.085 |
| 1 | 85 | 85 | -3.568 | -1.652 | -1.076 | -0.249 | 2.169 | -1.011 | 1.154 | 0.125 |
| 2 | 33 | 33 | -3.077 | -1.499 | -0.763 | 0.007 | 1.103 | -0.832 | 1.141 | 0.199 |
| 3 | 50 | 50 | -3.098 | -1.276 | 0.167 | 0.961 | 2.189 | -0.130 | 1.462 | 0.207 |

| **Analysis Variable : CTOT04_sig** | | | | | | | | | | |
| --- | --- | --- | --- | --- | --- | --- | --- | --- | --- | --- |
| **t Banff lesion score** | **N Obs** | **N** | **Minimum** | **Lower Quartile** | **Median** | **Upper Quartile** | **Maximum** | **Mean** | **Std Dev** | **Std Error** |
| 0 | 238 | 238 | -4.201 | -2.088 | -1.305 | -0.572 | 2.169 | -1.296 | 1.176 | 0.076 |
| 1 | 44 | 44 | -3.494 | -1.606 | -1.025 | -0.229 | 1.103 | -0.961 | 1.056 | 0.159 |
| 2 | 33 | 33 | -3.098 | -2.091 | -1.255 | 0.007 | 1.291 | -1.072 | 1.228 | 0.214 |
| 3 | 39 | 39 | -2.893 | -0.401 | 0.386 | 1.253 | 2.189 | 0.253 | 1.297 | 0.208 |

***The GLM Procedure***

| **Class Level Information** | | |
| --- | --- | --- |
| **Class** | **Levels** | **Values** |
| **g_lesions** | 4 | 0 1 2 3 |
| **ptc_lesions** | 4 | 0 1 2 3 |

| **Number of Observations Read** | 354 |
| --- | --- |
| **Number of Observations Used** | 354 |

| **Source** | **DF** | **Sum of Squares** | **Mean Square** | **F Value** | **Pr > F** |
| --- | --- | --- | --- | --- | --- |
| **Model** | 6 | 53.0422402 | 8.8403734 | 5.95 | <.0001 |
| **Error** | 347 | 515.6253352 | 1.4859520 |  |  |
| **Corrected Total** | 353 | 568.6675754 |  |  |  |

| **R-Square** | **Coeff Var** | **Root MSE** | **CTOT04_sig Mean** |
| --- | --- | --- | --- |
| 0.093275 | -114.6745 | 1.218996 | -1.063006 |

| **Source** | **DF** | **Type III SS** | **Mean Square** | **F Value** | **Pr > F** |
| --- | --- | --- | --- | --- | --- |
| **g_lesions** | 3 | 13.18021199 | 4.39340400 | 2.96 | 0.0325 |
| **ptc_lesions** | 3 | 11.91543242 | 3.97181081 | 2.67 | 0.0473 |

***The GLM Procedure***

| **Class Level Information** | | |
| --- | --- | --- |
| **Class** | **Levels** | **Values** |
| **i_lesions** | 4 | 0 1 2 3 |
| **t_lesions** | 4 | 0 1 2 3 |

| **Number of Observations Read** | 354 |
| --- | --- |
| **Number of Observations Used** | 354 |

| **Source** | **DF** | **Sum of Squares** | **Mean Square** | **F Value** | **Pr > F** |
| --- | --- | --- | --- | --- | --- |
| **Model** | 6 | 87.3994549 | 14.5665758 | 10.50 | <.0001 |
| **Error** | 347 | 481.2681206 | 1.3869398 |  |  |
| **Corrected Total** | 353 | 568.6675754 |  |  |  |

| **R-Square** | **Coeff Var** | **Root MSE** | **CTOT04_sig Mean** |
| --- | --- | --- | --- |
| 0.153692 | -110.7881 | 1.177684 | -1.063006 |

| **Source** | **DF** | **Type III SS** | **Mean Square** | **F Value** | **Pr > F** |
| --- | --- | --- | --- | --- | --- |
| **i_lesions** | 3 | 6.50717086 | 2.16905695 | 1.56 | 0.1979 |
| **t_lesions** | 3 | 23.37378241 | 7.79126080 | 5.62 | 0.0009 |

**SUPPLEMENTARY TABLE 5. Relationship of Composite Banff Scores by Urinary Cell**

**Three-Gene Signature Score in 354 Biopsy-Matched Urine Specimens**

**Banff g+ptc scores**

***The MEANS Procedure***

| **Analysis Variable : CTOT04_sig** | | | | | | | | | | |
| --- | --- | --- | --- | --- | --- | --- | --- | --- | --- | --- |
| **g+ptc Banff lesion scores** | **N Obs** | **N** | **Minimum** | **Lower Quartile** | **Median** | **Upper Quartile** | **Maximum** | **Mean** | **Std Dev** | **Std Error** |
| 0 | 143 | 143 | -4.201 | -2.237 | -1.362 | -0.619 | 1.494 | -1.418 | 1.184 | 0.099 |
| 1 | 63 | 63 | -4.118 | -2.230 | -1.431 | -0.466 | 2.189 | -1.185 | 1.327 | 0.167 |
| 2 | 41 | 41 | -3.077 | -2.164 | -1.179 | -0.324 | 1.291 | -1.171 | 1.067 | 0.167 |
| 3 | 48 | 48 | -3.086 | -1.591 | -0.594 | 0.489 | 1.932 | -0.536 | 1.285 | 0.185 |
| 4 | 20 | 20 | -3.568 | -1.420 | -0.283 | 0.363 | 1.093 | -0.574 | 1.301 | 0.291 |
| 5 | 19 | 19 | -1.440 | -0.872 | -0.572 | 0.684 | 2.169 | -0.171 | 0.989 | 0.227 |
| 6 | 20 | 20 | -2.937 | -1.167 | -0.482 | 0.214 | 2.107 | -0.520 | 1.235 | 0.276 |

**Banff i+t scores**

***The MEANS Procedure***

| **Analysis Variable : CTOT04_sig** | | | | | | | | | | |
| --- | --- | --- | --- | --- | --- | --- | --- | --- | --- | --- |
| **i+t Banff lesion scores** | **N Obs** | **N** | **Minimum** | **Lower Quartile** | **Median** | **Upper Quartile** | **Maximum** | **Mean** | **Std Dev** | **Std Error** |
| 0 | 184 | 184 | -4.201 | -2.223 | -1.346 | -0.589 | 1.505 | -1.366 | 1.150 | 0.085 |
| 1 | 52 | 52 | -3.568 | -1.896 | -1.197 | -0.503 | 2.169 | -1.143 | 1.274 | 0.177 |
| 2 | 36 | 36 | -2.701 | -1.444 | -1.001 | 0.053 | 1.014 | -0.797 | 1.003 | 0.167 |
| 3 | 9 | 9 | -2.230 | -1.893 | -1.631 | -0.948 | 1.103 | -1.309 | 1.031 | 0.344 |
| 4 | 21 | 21 | -3.077 | -2.179 | -0.910 | 0.007 | 0.784 | -1.086 | 1.199 | 0.262 |
| 5 | 19 | 19 | -3.098 | -1.649 | -0.401 | -0.035 | 1.291 | -0.685 | 1.118 | 0.257 |
| 6 | 33 | 33 | -2.893 | -0.275 | 0.663 | 1.289 | 2.189 | 0.327 | 1.395 | 0.243 |

**Banff g+ptc+i+t scores**

***The MEANS Procedure***

| **Analysis Variable : CTOT04_sig** | | | | | | | | | | |
| --- | --- | --- | --- | --- | --- | --- | --- | --- | --- | --- |
| **Total inflation (g+ptc+i+t)** | **N Obs** | **N** | **Minimum** | **Lower Quartile** | **Median** | **Upper Quartile** | **Maximum** | **Mean** | **Std Dev** | **Std Error** |
| 0 | 105 | 105 | -4.201 | -2.333 | -1.468 | -0.675 | 1.494 | -1.527 | 1.171 | 0.114 |
| 1 | 53 | 53 | -4.118 | -2.226 | -1.443 | -0.500 | 1.505 | -1.279 | 1.216 | 0.167 |
| 2 | 42 | 42 | -3.494 | -1.652 | -1.025 | -0.679 | 1.014 | -1.174 | 0.898 | 0.139 |
| 3 | 21 | 21 | -3.086 | -1.797 | -1.591 | -0.603 | 0.741 | -1.399 | 0.954 | 0.208 |
| 4 | 24 | 24 | -3.086 | -1.912 | -0.999 | 0.376 | 1.103 | -0.820 | 1.336 | 0.273 |
| 5 | 15 | 15 | -3.568 | -1.711 | -0.872 | 0.072 | 0.250 | -0.972 | 1.094 | 0.282 |
| 6 | 23 | 23 | -3.098 | -2.164 | -1.373 | -0.527 | 2.169 | -1.207 | 1.254 | 0.262 |
| 7 | 23 | 23 | -2.912 | -0.961 | -0.035 | 0.753 | 2.189 | -0.158 | 1.384 | 0.289 |
| 8 | 15 | 15 | -2.893 | -1.943 | -0.275 | 0.251 | 0.827 | -0.763 | 1.205 | 0.311 |
| 9 | 19 | 19 | -2.091 | -0.642 | 0.663 | 1.253 | 1.932 | 0.358 | 1.127 | 0.259 |
| 10 | 6 | 6 | -0.523 | -0.512 | -0.206 | 0.283 | 1.093 | -0.012 | 0.639 | 0.261 |
| 11 | 5 | 5 | -1.649 | -1.394 | -0.742 | 0.386 | 1.289 | -0.422 | 1.238 | 0.554 |
| 12 | 3 | 3 | 0.177 | 0.177 | 0.718 | 2.107 | 2.107 | 1.001 | 0.996 | 0.575 |

***The GLM Procedure***

| **Class Level Information** | | |
| --- | --- | --- |
| **Class** | **Levels** | **Values** |
| **g_ptc_** | 7 | 0 1 2 3 4 5 6 |
| **i_t_** | 7 | 0 1 2 3 4 5 6 |

| **Number of Observations Read** | 354 |
| --- | --- |
| **Number of Observations Used** | 354 |

| **Source** | **DF** | **Sum of Squares** | **Mean Square** | **F Value** | **Pr > F** |
| --- | --- | --- | --- | --- | --- |
| **Model** | 12 | 110.2092415 | 9.1841035 | 6.83 | <.0001 |
| **Error** | 341 | 458.4583339 | 1.3444526 |  |  |
| **Corrected Total** | 353 | 568.6675754 |  |  |  |

| **R-Square** | **Coeff Var** | **Root MSE** | **CTOT04_sig Mean** |
| --- | --- | --- | --- |
| 0.193803 | -109.0780 | 1.159505 | -1.063006 |

| **Source** | **DF** | **Type III SS** | **Mean Square** | **F Value** | **Pr > F** |
| --- | --- | --- | --- | --- | --- |
| **g_ptc_** | 6 | 23.45829562 | 3.90971594 | 2.91 | 0.0089 |
| **i_t_** | 6 | 51.68647628 | 8.61441271 | 6.41 | <.0001 |

| **Parameter** | **Estimate** |  | **Standard Error** | **t Value** | **Pr > \|t\|** |
| --- | --- | --- | --- | --- | --- |
| **Intercept** | 0.511081472 | B | 0.32307348 | 1.58 | 0.1146 |
| **g_ptc_ 0** | -0.588947559 | B | 0.28876011 | -2.04 | 0.0422 |
| **g_ptc_ 1** | -0.481076441 | B | 0.30344498 | -1.59 | 0.1138 |
| **g_ptc_ 2** | -0.553000878 | B | 0.32139564 | -1.72 | 0.0862 |
| **g_ptc_ 3** | -0.093064856 | B | 0.31307564 | -0.30 | 0.7664 |
| **g_ptc_ 4** | -0.078714115 | B | 0.37209817 | -0.21 | 0.8326 |
| **g_ptc_ 5** | 0.368384086 | B | 0.37327526 | 0.99 | 0.3244 |
| **g_ptc_ 6** | 0.000000000 | B | . | . | . |
| **i_t_ 0** | -1.398090304 | B | 0.23682698 | -5.90 | <.0001 |
| **i_t_ 1** | -1.308103487 | B | 0.26743182 | -4.89 | <.0001 |
| **i_t_ 2** | -1.019626609 | B | 0.28607599 | -3.56 | 0.0004 |
| **i_t_ 3** | -1.499356079 | B | 0.44214310 | -3.39 | 0.0008 |
| **i_t_ 4** | -1.343014971 | B | 0.32824530 | -4.09 | <.0001 |
| **i_t_ 5** | -0.911071491 | B | 0.33703745 | -2.70 | 0.0072 |
| **i_t_ 6** | 0.000000000 | B | . | . | . |

| **Note:** | The X'X matrix has been found to be singular, and a generalized inverse was used to solve the normal equations. Terms whose estimates are followed by the letter 'B' are not uniquely estimable. |
| --- | --- |
